# FAK and p130Cas modulate stiffness-mediated early transcription and cellular metabolism

**DOI:** 10.1101/2024.01.15.575789

**Authors:** Bat-Ider Tumenbayar, Vincent M. Tutino, Joseph A. Brazzo, Peng Yao, Yongho Bae

**Affiliations:** Department of Pharmacology and Toxicology, Jacobs School of Medicine and Biomedical Sciences, University at Buffalo, Buffalo, NY 14203, USA; Department of Pathology and Anatomical Sciences, Jacobs School of Medicine and Biomedical Sciences, University at Buffalo, Buffalo, NY 14203, USA; Department of Biomedical Engineering, School of Engineering and Applied Sciences, University at Buffalo, Buffalo, NY 14260, USA; Department of Neurosurgery, Jacobs School of Medicine and Biomedical Sciences, University at Buffalo, Buffalo, NY 14203, USA; Aab Cardiovascular Research Institute, Department of Medicine, University of Rochester School of Medicine and Dentistry, Rochester, NY 14642, USA

## Abstract

Cellular metabolism is influenced by the stiffness of the extracellular matrix. Focal adhesion kinase (FAK) and its binding partner, p130Cas, transmit biomechanical signals about substrate stiffness to the cell to regulate a variety of cellular responses, but their roles in early transcriptional and metabolic responses remain largely unexplored. We cultured mouse embryonic fibroblasts with or without siRNA-mediated FAK or p130Cas knockdown and assessed the early transcriptional responses of these cells to placement on soft and stiff substrates by RNA sequencing and bioinformatics analyses. Exposure to the stiff ECM altered the expression of genes important for metabolic and biosynthetic processes, and these responses were influenced by knockdown of FAK and p130Cas. Our findings reveal that FAK-p130Cas signaling mechanotransduces ECM stiffness to early transcriptional changes that alter cellular metabolism and biosynthesis.

## BACKGROUND

Extracellular matrix (ECM) is part of a dynamic microenvironment that modulates various energy-demanding cellular processes, including cell adhesion, spreading, differentiation, migration, and proliferation [1, 2, 3, 4, 5]. There is evidence that the stiffness of the ECM serves as a biomechanical signal that influences mitochondrial network structure and function [6, 7, 8]. For example, a stiff ECM signals mitochondrial reprogramming to increase energy production for adhesion of the cell to a stiffer substrate [9]. This reprogramming and associated changes to a variety of cell processes involve transcriptional changes as the cells adapt to their microenvironment. The mechanism directing these broad changes is not clearly understood.

The mechanotransduction of ECM stiffness begins with the direct interaction between ECM components and integrins, which are cell adhesion receptors and non-catalytic proteins that regulate critical mediators of cell metabolism [10]. Integrins signal through their association with cytosolic proteins, including a non-receptor tyrosine kinase known as focal adhesion kinase (FAK) [11, 12, 13]. Our previous studies showed that a stiffer ECM leads to increased phosphorylation of FAK and one of its binding partners, p130Cas [14, 15, 16]. We identified p130Cas as a target of FAK because FAK inhibition reduced stiffness-induced p130Cas phosphorylation. This stiffness-mediated activation of FAK and p130Cas regulates several cellular processes, including cell cycle progression, proliferation [17, 18, 19], cell adhesion, and motility [20, 21, 22] by signaling through Rac, ERK, and AKT pathways. Thus, FAK and p130Cas are positioned to act as mechanotransducers, signaling that adaptations to cell processes are needed to adapt to the change in the microenvironment.

To investigate whether FAK and p130Cas modulate transcriptional programming in response to ECM stiffness, we used next-generation sequencing and a comprehensive bioinformatics approach to compare the transcriptomes of mouse embryonic fibroblasts (MEFs) with and without FAK and p130Cas knockdown and the global transcriptional differences between cells on soft and stiff substrates. We mapped out protein–protein interactions among commonly regulated genes to reveal which metabolic and biosynthetic processes are modulated by FAK-p130Cas signaling as cells adapt to physical changes in their microenvironment.

## METHODS

### Cell culture

Spontaneously immortalized mouse embryonic fibroblasts (MEFs), provided by the Assoian Laboratory at the University of Pennsylvania, were cultured in low-glucose Dulbecco’s Modified Eagle’s Medium (DMEM; cat. no. 10-014-CV, Corning) supplemented with 50 μg/ml gentamicin solution (cat. no. 30-005-CR, Corning) and 10% fetal bovine serum (FBS; cat. no. F2442, Sigma-Aldrich). Before seeding MEFs on fibronectin (cat. no. 341631, Calbiochem)-coated soft or stiff polyacrylamide hydrogels [23] for experimentation, cells were synchronized in the G_0_ cell cycle phase, in which cells near confluency (∼80%) were serum starved by incubation for 24 h in DMEM with 1 mg/ml heat-inactivated, fatty-acid-free bovine serum albumin (BSA; cat. no. 5217, Tocris). Subsequently, serum-starved MEFs were replated for 1 h on polyacrylamide hydrogels with 10% FBS, and total lysates were collected for immunoblotting or RNA sequencing analyses.

### siRNA transfection

MEFs were transfected with 200 nM FAK or p130Cas siRNAs using Lipofectamine 2000 reagent (cat. no. 11668019, Invitrogen) in Opti-MEM (cat. no. 31985070, Gibco) according to previously described methods [4, 14, 16]. MEFs were transfected with siRNA for 5 h followed by immediate serum starvation in DMEM with 1 mg/ml BSA for 24 h. Cells were then trypsinized and seeded on fibronectin-coated hydrogels with DMEM containing 10% FBS. A non-targeting siRNA (cat. no. AM4611, Ambion) served as an experimental control. FAK and p130Cas siRNAs were obtained from Ambion: FAK siRNA #1 (ID: 157448): CCUAGCAGACUUUAACCAAtt; FAK siRNA #2 (ID: 61352): GGCAUGGAGAUGCUACUGAtt; p130Cas siRNA #1 (ID: 161328): GCCAAUCGGCAUCUUCCUUtt; p130Cas siRNA #2 (ID: 161329): GCUGAAACAGUUUGAGCGAtt.

### Immunoblotting

As previously described [4, 14, 16], total cell lysates were collected from MEFs cultured on polyacrylamide hydrogels by incubating the hydrogels face down for 1 to 2 min at room temperature on 5× SDS sample buffer containing β-mercaptoethanol. Equal amounts of extracted protein were fractionated on reducing 10% SDS-polyacrylamide gels, and the fractioned proteins were subsequently transferred electrophoretically onto nitrocellulose blotting membranes. These membranes were probed with antibodies against FAK (cat. no. 610087, BD Transduction), p130Cas (cat. no. 610271, BD Transduction), or GAPDH (cat. no. sc-25778, Santa Cruz Biotechnology). Immunoblot signals were detected using enhanced chemiluminescence.

### RNA sample preparation for RNA-Seq analysis

RNA sample preparation and RNA-Seq analysis were in accordance with established protocols, as described in our previous research [3]. Triplicate samples were generated for MEFs cultured under four different conditions: (i) soft hydrogels with non-targeting control siRNA, (ii) stiff hydrogels with non-targeting control siRNA, (ii) stiff hydrogels with FAK siRNA #1, and (iv) stiff hydrogels with p130Cas siRNA #2. Total RNA was extracted using TRIzol reagent (cat. no. 15596018, Invitrogen) and subsequently purified with an RNeasy kit (cat. no. 74106, Qiagen). Libraries were prepared using the Illumina TruSeq Stranded total RNA kit with the Ribo-Zero plus rRNA depletion kit, followed by sequencing on an Illumina HiSeq 4000 PE100 sequencer. Sequencing coverage was approximately 40 million reads per sample.

### RNA-Seq data preprocessing and alignment

To evaluate the quality of RNA-Seq, we performed quality control analysis using both FastQC before alignment and MultiQC after alignment. The sequencing quality was assessed using FastQC version 0.11.9 [24], and potential contamination was detected by using FastQ Screen version 0.14.1 [25]. Summaries of FastQC and FastQ Screen quality reports were generated using MultiQC version 1.9 [26]. No adapter trimming was performed. Genomic alignments were performed using HISAT2 version 2.2.1 [27] with default parameters. The UCSC mm10 reference was used for the reference genome and gene annotation set. Sequence alignments were compressed and sorted into binary alignment map (BAM) files using SAMtools version 1.3. Mapped reads for genomic features were counted using Subread featureCounts version 2.0.0 [28] using the parameters -s 2 –g gene_name –t exon –Q 60 -B -C; the annotation file specified with –a was the UCSC mm10 reference provided by Illumina’s iGenomes. After the counting, expression values for 23,418 transcripts were defined. Alignment statistics and feature assignment statistics were once again summarized using MultiQC.

### Identification of differentially expressed genes

Data were first preprocessed to ensure the quality of the datasets. Gene entries with expression in all samples were retained for differential expression analysis. We employed ggplot2 [29] and pheatmap [30] packages to create visualizations, including a principal-component analysis plot and a sample correlation heat map, using a normalized count dataset. The DESeq2 [31] package in R was applied to identify differentially expressed genes (DEGs) between the following groups: stiff hydrogels with FAK siRNA, stiff hydrogels with p130Cas siRNA, soft hydrogels with control siRNA, and stiff hydrogels with control siRNA. DEGs were selected on the basis of the following thresholds as recently described in [3]: Benjamini-Hochberg adjusted *p* value of <0.05, absolute log_2_(fold-change) of >0.32, and baseMean intensity of >500. Python’s bioinfokit [32] and R’s pheatmap packages [30] were used to illustrate gene expression patterns and distribution.

### Functional enrichment analysis

To gain a deeper understanding of the biological processes associated with DEGs, a gene enrichment analysis was performed to examine biological processes and Kyoto Encyclopedia of Genes and Genomes (KEGG) pathways. The analysis was carried out using the g:GOSt function on the gProfiler web server (https://biit.cs.ut.ee/gprofiler/gost) [33]. The significance threshold was set to the Benjamini-Hochberg FDR (false discovery rate) value, and significant GO terms and KEGG pathways were defined by an adjusted *p* value of ≤0.05. For visualization purposes, bubble plots representing the top 20 enriched GO terms and KEGG pathways were generated using the SRplot online server.

### Ingenuity Pathway Analysis

Commonly regulated DEGs from the FAK siRNA and p130Cas siRNA datasets were uploaded to the Ingenuity Pathway Analysis (IPA) software [34], using the expression log ratio and adjusted *p* values from the FAK dataset as the observation. IPA’s Core Analysis function was used to gain a deeper understanding of altered signaling pathways in response to FAK and p130Cas knockdown. The Diseases and Functions and Pathways features were used to determine significantly affected pathways and diseases [absolute activation z score ≥ 2; -log_10_(*p* value) ≥ 2] on the basis of common molecules. The Network Analysis feature was used to explore transcriptional networks leading to metabolic processes. The statistical values for Network Analysis were computed based on the p-score, derived from *p* values and -log_10_(*p* values). Additionally, the “My pathway” tool was used to illustrate known relationships between molecules or molecules to functions.

### GO term analysis and protein–protein interaction network

We performed GO term “biological process” enrichment analysis on DEGs commonly regulated in FAK siRNA and p130Cas siRNA datasets and visualized the results using the ClueGO plugin within Cytoscape [35]. The analysis used a two-sided hypergeometric test with Benjamini-Hochberg correction (*p* < 0.05) and a kappa score threshold of 0.5 to determine GO term network connectivity. The nodal network representation for GO biological process terms was constrained (*p* < 0.0001) and global network specificity to fit the terms within the graphic. All significantly enriched GO biological processes (Benjamini Hochberg corrected p value < 0.05) were included in the Supplementary Table S2. To explore protein–protein interactions (PPIs) among DEGs, the STRING website [36] was used to construct the PPI network. STRING website’s k-means clustering online tool was used to identify clusters within the PPI network. Orphan and non-present protein entries in the dataset were filtered and the network was visualized in Cytoscape. FAK siRNA expression data were incorporated into the node table, with log₂(fold-change) values indicating expression levels and node color representing intensity. Additionally, we identified the top 10 hub genes using the Density of Maximum Neighborhood Component (DMNC) topological algorithm within the cytoHubba application [37].

### RNA binding protein motif enrichment analysis

The Transite transcript set motif analysis online tool [38] was used to identify enriched RNA-binding protein (RBP) motifs within the commonly regulated genes affected by FAK and p130Cas knockdown. Separate analyses were conducted on the 5’ and 3’ untranslated regions (UTRs), employing a matrix-based transcript set motif analysis, with the whole transcript list as the background. The analysis pipeline was configured with the Benjamini-Hochberg method, allowing for a maximum of 50 binding sites per mRNA. RBP motifs with a *p* value of <0.05 were considered significantly enriched and hierarchically clustered using Euclidean distance and the ward.D2 clustering method. The resulting dendrogram displaying differentially represented RBP motifs was visualized using the ggdendro package in R. The PPI network of all enriched RBPs was visualized using Cytoscape [39]. The STRING enrichment application [36] was applied to perform enrichment analysis for RBPs and proteins correlated to the following significantly enriched biological process GO categories: regulation of mRNA metabolic process (GO:1903311), gene expression (GO:0010467), regulation of RNA metabolic process (GO:0051252), mRNA processing (GO:0006397), RNA splicing (GO:0008380), regulation of translation (GO:0006417), macromolecule metabolic process (GO:0043170), cellular nitrogen compound metabolic process (GO:0034641), regulation of cellular metabolic process (GO:0031323), and regulation of primary metabolic process (GO:0080090).

### Statistical analysis

Statistical significance was assessed for the data in Figure 1B using paired, one-tailed Student’s *t* tests. The graphs present means + SD from the indicated number of independent experiments.

**Figure 1.**
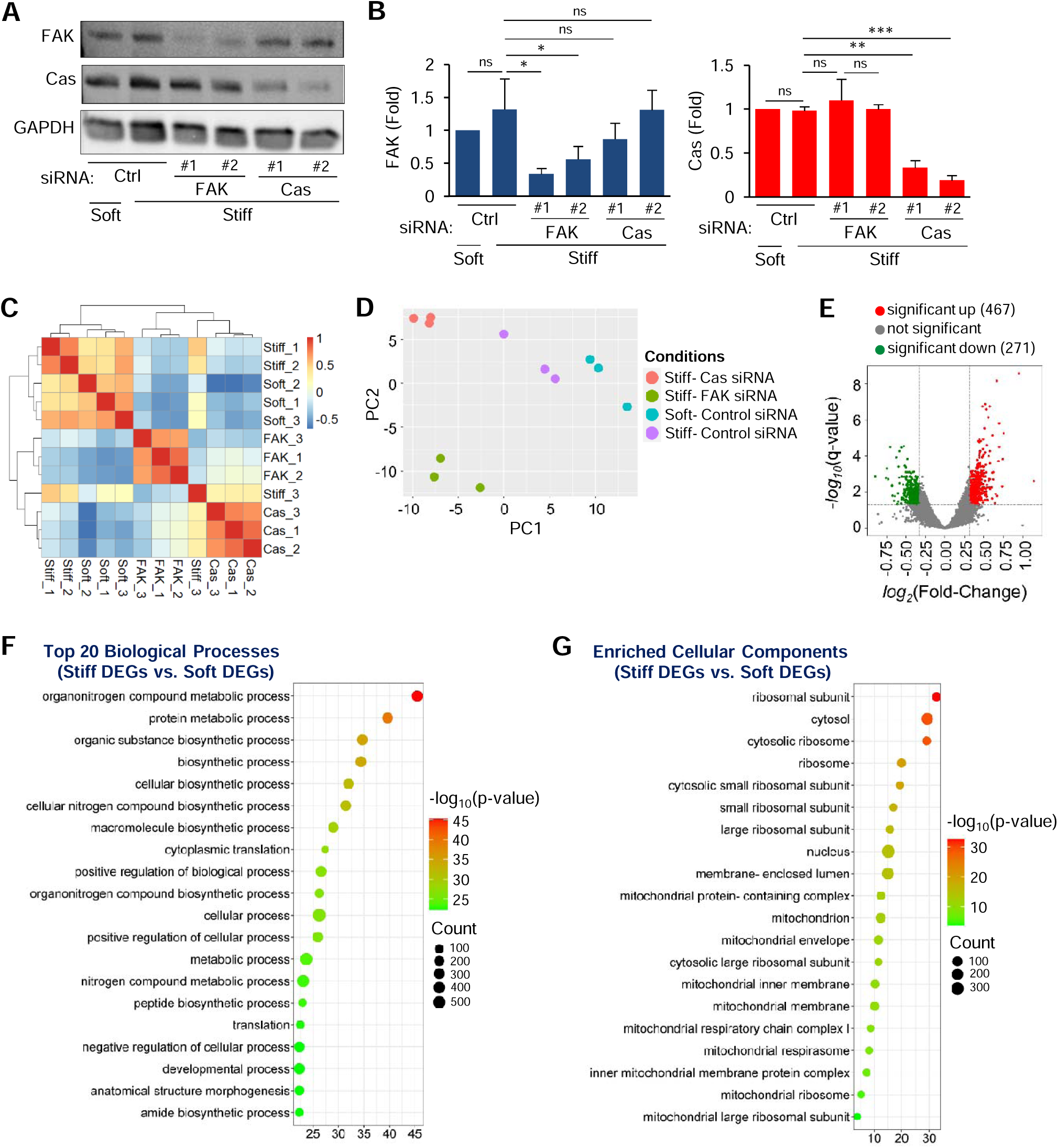
RNA-Seq analysis identify stiffness-induced early transcriptional changes and associated biological processes and cellular components. Mouse embryonic fibroblasts (MEFs) were transfected with non-targeting control siRNA or siRNAs targeting FAK and p130Cas, synchronized to G_0_, and plated on fibronectin-coated soft or stiff substrates for 1 h. (A) Immunoblots of total cell lysates showing protein levels of FAK and p130Cas with GAPDH as the loading control. (B) Graphs present mean + SD, normalized to GAPDH abundance, and plotted relative to the signal on the soft hydrogels, *n* = 3. \**p* < 0.05, \*\**p* < 0.01, \*\*\**p* < 0.001; ns, not significant. Correlation heat map (C) and principal-component analysis plot (D) for the entire transcriptome list. (E) Volcano plot illustrates the distribution of differentially expressed genes (DEGs) in response to stiffer ECM; significance determined by adjusted *p* value of <0.05; log_2_(fold-change) <-0.32 or >0.32; and baseMean > 500. Bubble plots depict the top 20 significantly enriched biological processes (F) and cellular components (G) among DEGs in response to stiff versus soft substrate.

## RESULTS

### Stiff ECM influences metabolic and biosynthetic processes

To investigate the roles of substrate stiffness and FAK/p130Cas in the early transcriptional response, a whole-transcriptome analysis was performed using mRNA samples from MEFs transfected with FAK siRNA, p130Cas siRNA, or non-targeting control siRNA and cultured for 1 h on soft (3–8 kPa) or stiff (18–24 kPa) [4] polyacrylamide-based hydrogels. Immunoblotting analysis of total lysates confirmed that the siRNAs successfully downregulated their targets (**Fig. 1A-B**). RNA sequencing (RNA-Seq) analysis of mRNAs isolated from these samples yielded an average of 56.3 million sequences per sample with a 74.9% read mapping rate (**Table S1**). Expression values of 23,418 high-quality transcripts were obtained by counting mapped reads for genomic features. The relationships between samples according to gene expression were assessed in an unsupervised manner and can be visualized in a correlation heat map (**Fig. 1C**) and principal-component analysis plot (**Fig. 1D**). We then compared high stiffness versus low stiffness and identified 738 differentially expressed genes (DEGs): 467 were upregulated and 271 were downregulated (**Fig. 1E**). To better understand this early transcriptional response to ECM stiffness, we performed a Gene Ontology (GO) enrichment analysis. The top biological process categories enriched among the DEGs were mainly related to metabolic and biosynthetic processes, including “organonitrogen compound metabolic process,” “protein metabolic process,” “organic substance biosynthetic process,” “biosynthetic process,” and “cellular biosynthetic process” (**Fig. 1F**). Additionally, the analysis of cellular component categories indicated significant enrichment for cellular metabolism, including “ribosomal subunit,” “ribosome,” “cytosolic ribosome,” “mitochondrial protein-containing complex,” and “inner mitochondrial membrane protein complex,” in response ECM stiffness (**Fig. 1G**). These data show that ECM stiffness substantially impacts the cellular transcriptome, particularly influencing processes related to metabolism and biosynthesis.

### FAK modulates stiffness-mediated metabolic and biosynthesis processes

To examine the role of FAK in the early transcriptional response to ECM stiffness, we conducted a differential expression analysis using DESeq2 on RNA-Seq data obtained from MEFs transfected with FAK or non-targeting control siRNAs and cultured on stiff hydrogels, with triplicate samples for each condition. Genes were filtered by significance thresholds (see methods and reference [3]), resulting in the identification of 924 DEGs: 385 showing upregulation and 539 displaying downregulation. The distribution and expression patterns of these DEGs are displayed in a volcano plot (**Fig. 2A**) and correlation heat map (**Fig. 2B**). To further investigate biological functions associated with DEGs, we conducted GO enrichment analyses. Biological process categories that were enriched among the downregulated DEGs in response to FAK knockdown included “developmental process,” “regulation of response to stimuli,” “organonitrogen compound metabolic process,” and “protein metabolic process” (**Fig. 2C**). Those enriched among upregulated DEGs included “regulation of biological processes,” “developmental process,” “positive regulation of metabolic processes,” “cell differentiation,” and “regulation of cell signaling” (**Fig. 2D**). The KEGG pathway enrichment analysis found enrichment in pathways such as “MAPK signaling pathway,” “FoxO signaling pathway,” “pathways in cancer,” and “biosynthesis of amino acids” (**Fig. 2E**). An Ingenuity Pathway Analysis (IPA) predicted significant and differential changes with positive enrichment of five canonical pathways (including “superpathway of cholesterol biosynthesis” and “cell cycle control of chromosomal replication”) and negative enrichment of seven canonical pathways (including “pulmonary fibrosis idiopathic signaling pathway” and “GP6 signaling pathway”) in response to FAK knockdown (**Fig. 2F**).

**Figure 2.**
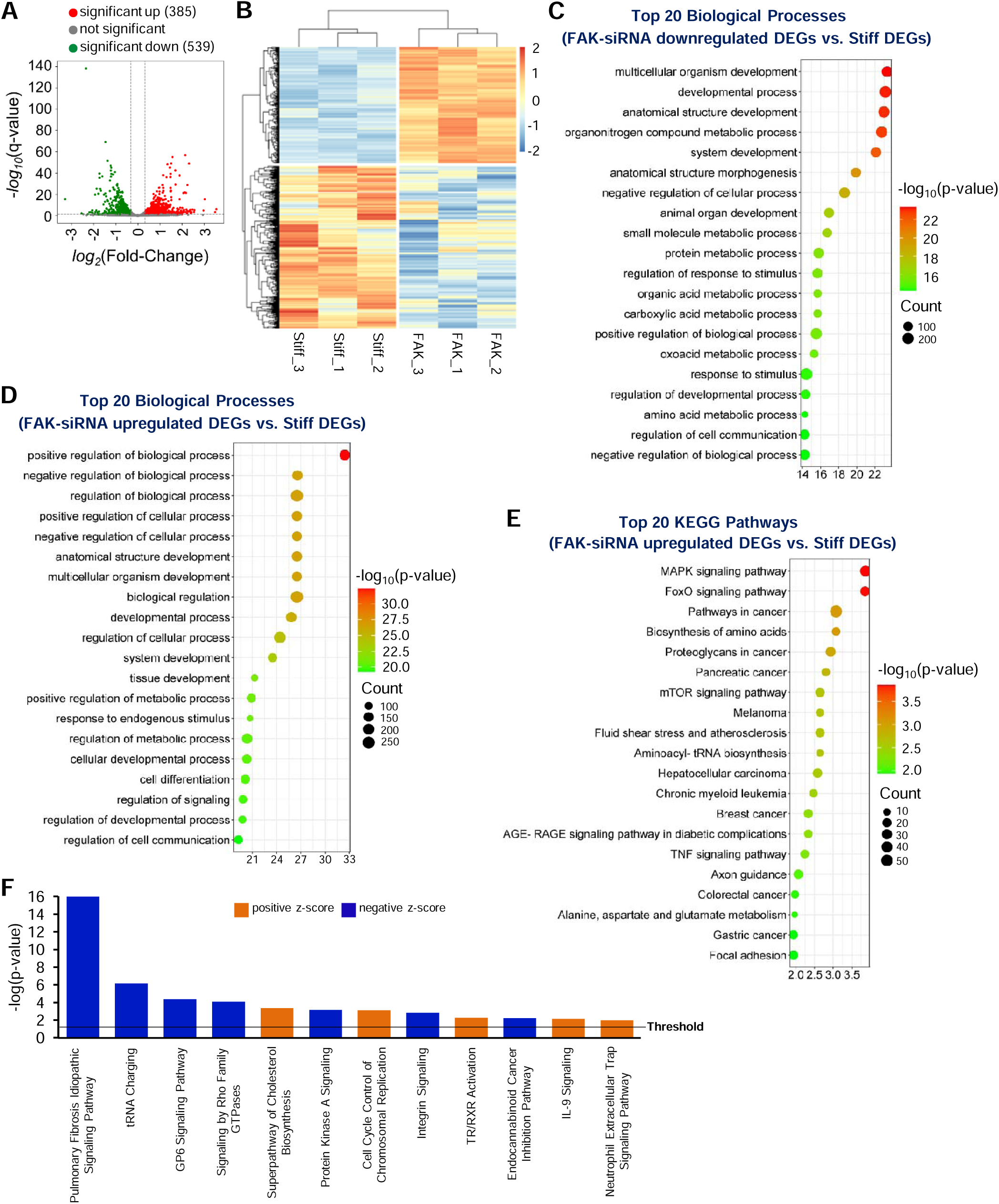
FAK knockdown affects stiffness-mediated cellular function and metabolic processes. Volcano plot (**A**) and heat map (**B**) to visualize the distribution and expression patterns, respectively, of differentially expressed genes (DEGs) in response to FAK knockdown. Bubble plots show the top 20 enriched biological processes for significantly downregulated (**C**) and upregulated (**D**) DEGs comparing FAK siRNA to control siRNA in cells on stiff hydrogels. (**E**) Top 20 enriched KEGG pathways for DEGs with FAK siRNA versus control siRNA in cells on stiff hydrogels. (**F**) Histogram represents significantly and differentially activated or inhibited [absolute activation z score ≥ 2; -log_10_(*p* value) ≥ 2)] canonical pathways in response to FAK knockdown.

We further compared data from FAK siRNA-transfected cells on stiff hydrogels with data from control siRNA-treated cells on soft hydrogels, identifying 1,929 DEGs (924 upregulated and 1,005 downregulated) (**Fig. S1A, B**). The enrichment analysis of downregulated DEGs revealed predominantly metabolic and biosynthetic processes influenced by FAK knockdown, including “organonitrogen compound metabolic process,” “organonitrogen compound biosynthesis process,” “protein metabolic process,” and “peptide biosynthetic process” (**Fig. S1C**); upregulated DEGs showed enrichment in biological processes such as “multicellular organism development,” “positive regulation of biological process,” “regulation of primary metabolic process,” and “anatomical structure development” (**Fig. S1D**). The KEGG pathway analysis identified enrichment in pathways including “biosynthesis of amino acids,” “carbon metabolism,” “VEGF signaling pathway,” “metabolic pathways,” and “FoxO signaling pathway” among others (**Fig. S1E**). Furthermore, IPA canonical pathway analysis predicted significant and differential changes in 111 canonical pathways, including positive enrichment of 68 pathways (including “mitochondrial dysfunction,” “ERK/MAPK signaling,” and “Rho GTPase cycle”) and negative enrichment of 43 pathways (including “selenoamino acid metabolism” and “mitochondrial translation”) (see **Fig. S1F** for a histogram of the top 20). These findings indicate that FAK’s role in the early transcriptional response to ECM stiffness involves modulation of critical cell functions, including metabolism and biosynthesis.

### p130Cas predominantly impacts stiffness-mediated metabolic and biosynthetic processes

To assess the impact of p130Cas knockdown on the early transcriptional response to ECM stiffness, we compared RNA-Seq data from MEFs treated with p130Cas siRNA with data from those treated with non-targeting control siRNA, both cultured on stiff hydrogels. We identified 952 DEGs (453 upregulated DEGs and 499 downregulated DEGs), which are presented in a volcano plot (**Fig. 3A**) and a heat map (**Fig. 3B**). Further functional enrichment analysis identified numerous metabolic and biosynthetic processes, such as “organonitrogen compound metabolic process,” “small molecule metabolic process,” “organic substance biosynthetic process,” and “primary metabolic process,” that were enriched among downregulated DEGs (**Fig. 3C**). Additionally, biological processes such as “anatomical structure development,” “developmental process,” “regulation of cellular response,” and “regulation of signaling” were enriched among upregulated DEGs (**Fig. 3D**). KEGG pathway analysis of all DEGs indicated enrichment in various disease pathways, such as “prion disease,” “Alzheimer disease,” “Parkinson disease,” and “Huntington disease.” There was also enrichment of “metabolic pathways,” “steroid biosynthesis,” “carbon metabolism,” and “valine, leucine, and isoleucine degradation pathways” (**Fig. 3E**). Moreover, IPA canonical pathway analysis predicted positive enrichment of 12 canonical pathways (including “pulmonary fibrosis idiopathic signaling pathway” and “superpathway of cholesterol biosynthesis”) and negative enrichment of 6 pathways (including “microautophagy signaling pathway” and “oxidative phosphorylation”) in response to p130Cas knockdown (**Fig. 3F**).

**Figure 3.**
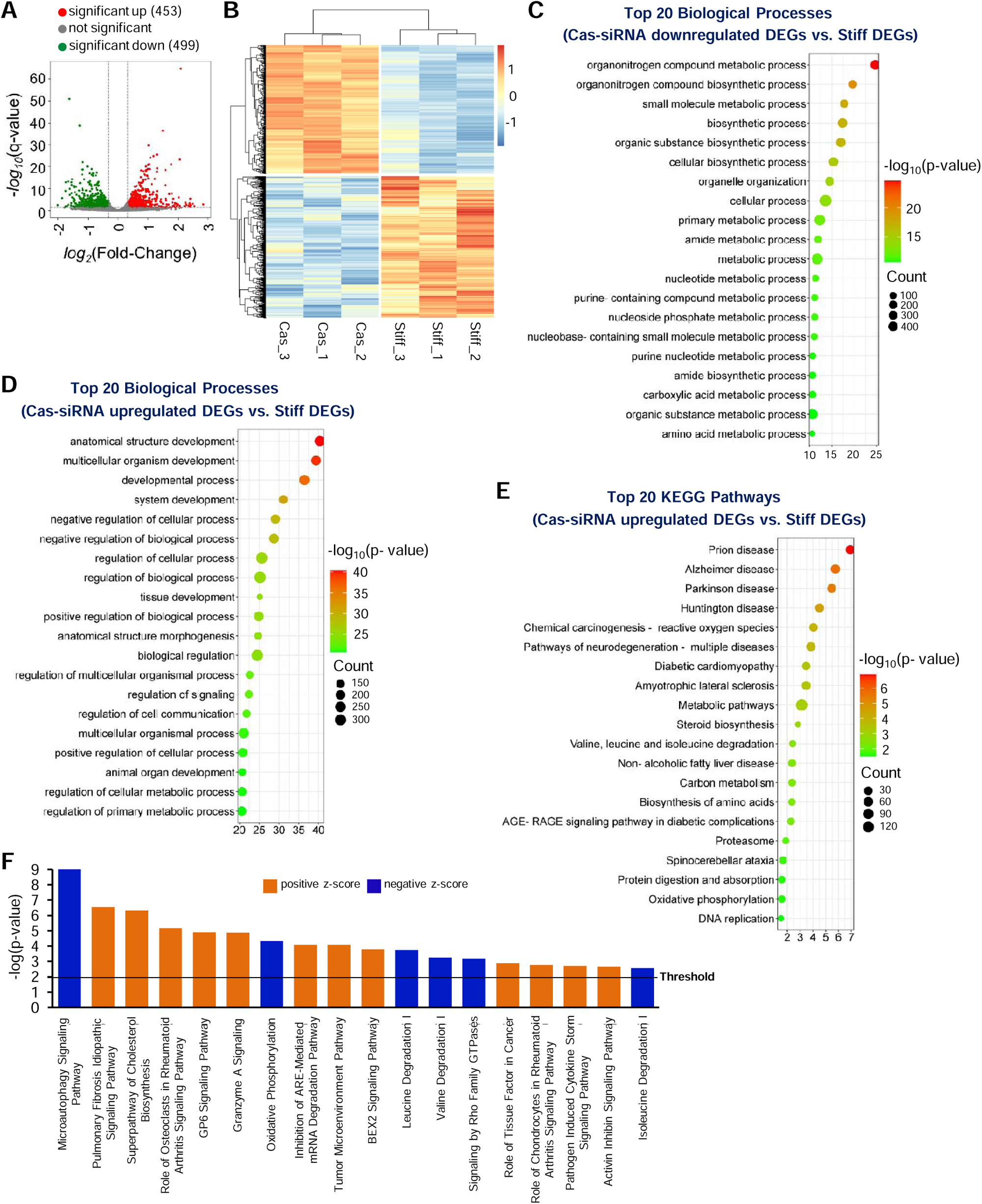
p130Cas knockdown results in significant changes in metabolic and biosynthetic processes. Volcano plot (**A**) and heat map (**B**) to visualize the distribution and expression patterns, respectively, of differentially expressed genes (DEGs) in response to p130Cas knockdown. Bubble plots show the top 20 enriched biological processes for significantly downregulated (**C**) and upregulated (**D**) DEGs comparing p130Cas siRNA to control siRNA in cells on stiff hydrogels. (**E**) Top 20 enriched KEGG pathways for DEGs with p130Cas siRNA versus control siRNA in cells on stiff hydrogels. (**F**) Histogram represents significantly and differentially activated or inhibited [absolute activation z score ≥ 2; -log_10_(*p* value) ≥ 2)] canonical pathways in response to p130Cas knockdown.

We also compared the data from cells transfected with p130Cas siRNA and cultured on stiff hydrogels to data from cells treated with control siRNA and cultured on soft hydrogels. This analysis revealed 2,705 DEGs: 1,367 upregulated and 1,338 downregulated (**Fig. S2A, B**). The enrichment analysis of these downregulated DEGs unveiled mostly metabolic and biosynthetic processes affected by p130Cas knockdown, including “organonitrogen compound biosynthetic process,” “organonitrogen compound metabolic process,” “peptide biosynthetic process,” and “protein metabolic process” (**Fig. S2C**). Additionally, upregulated DEGs in cells with p130Cas knockdown were associated with biological processes such as “anatomical structure development,” “multicellular organism development,” “developmental process,” “negative regulation of cellular process,” and “negative regulation of biological process” (**Fig. S2D**). The KEGG pathway analysis in this dataset highlighted disease pathways such as “Huntington disease,” “prion disease,” “Parkinson disease,” and “Alzheimer’s disease” as well as “oxidative phosphorylation” and “metabolic pathways” (**Fig. S2E**). Moreover, IPA canonical pathway analysis predicted significant and differential changes in 150 canonical pathways, including positive enrichment of 86 pathways (including “mitochondrial dysfunction,” “granzyme A signaling, and “Rho GTPase cycle”) and negative enrichment of 64 pathways (including “selenoamino acid metabolism,” “electron transport, ATP synthesis, and heat production by uncoupling proteins,” “oxidative phosphorylation,” and “mitochondrial translation”) (see **Fig. S2F** for a histogram of the top 20). The impact of p130Cas knockdown on biological processes and KEGG pathways was comparable to that observed with FAK knockdown, further emphasizing the roles of p130Cas and FAK in cellular metabolism, biosynthesis, and disease-related mechanisms.

### FAK-p130Cas signaling affects metabolic processes and RNA splicing

We further analyzed our dataset to identify DEGs in MEFs cultured on stiff hydrogels that were commonly regulated with FAK and p130Cas knockdown: 151 DEGs were commonly downregulated (**Fig. 4A**) and 157 DEGs were commonly upregulated (**Fig. 4B**) by FAK and p130Cas knockdown. GO analysis of these genes showed they were primarily enriched in “metabolic processes,” “tissue development,” “cell differentiation,” “regulation of signal transduction,” “apoptosis,” and “response to external stimulus” (**Fig. 4C**) (see **Table S2** for all enriched biological processes).

**Figure 4.**
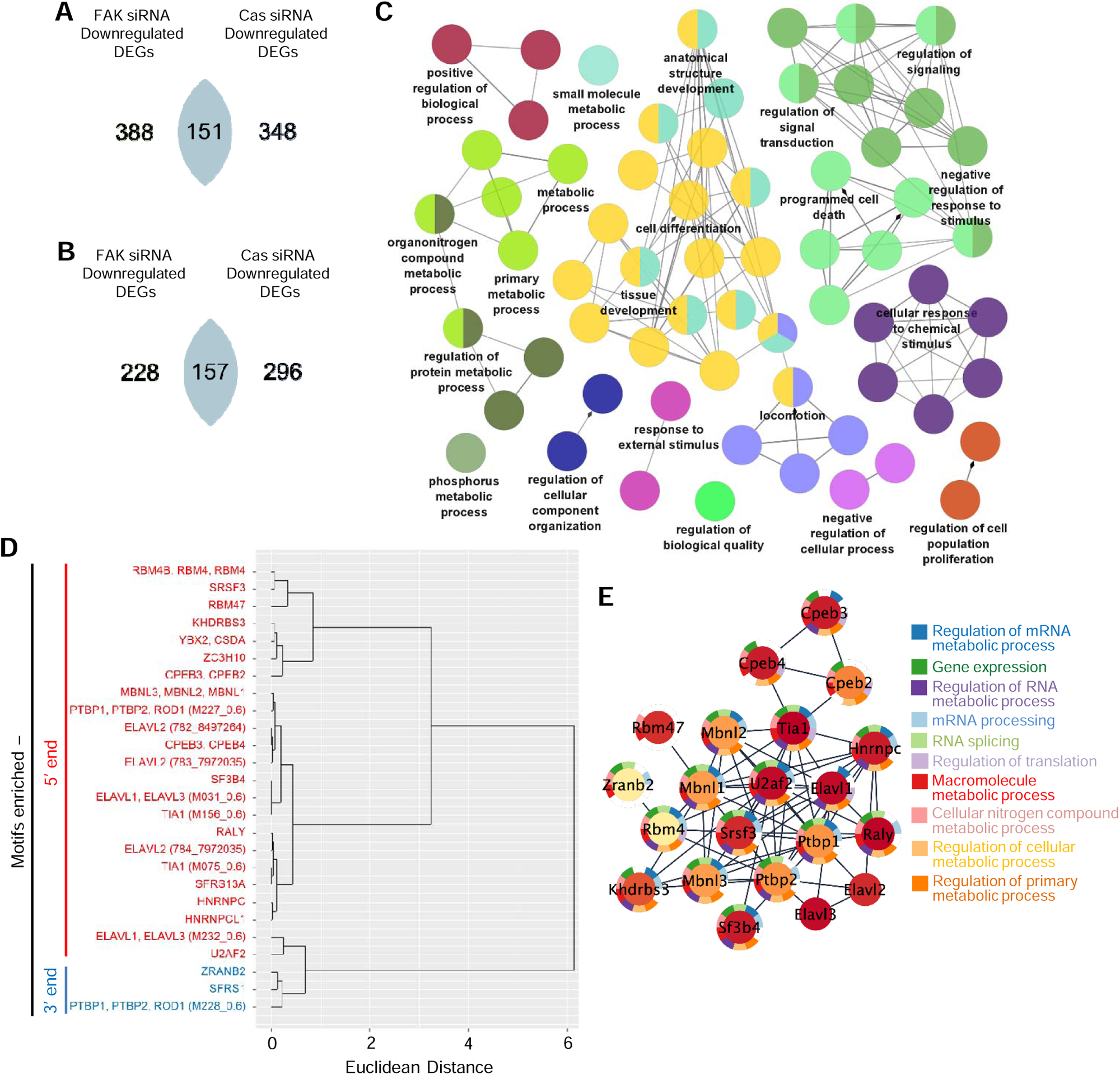
Metabolic processes are impacted by commonly regulated DEGs influenced by FAK-p130Cas signaling. Venn diagrams show commonly downregulated (**A**) and upregulated (**B**) differentially expressed genes (DEGs) in cells on stiff hydrogels with FAK siRNA or p130Cas siRNA. (**C**) Gene ontology (GO) analysis identifies a network of significantly connected and grouped biological processes (*p*□<□0.0001 by two-sided hypergeometric test with Benjamini– Hochberg correction; global network specificity) among common DEGs of FAK-p130Cas signaling. (**D**) Dendrogram shows hierarchical clustering of significantly enriched (*p* < 0.05) RNA-binding protein motifs identified by the Transite set motif analysis tool in 3′ and 5′ untranslated regions of common DEGs. (**E**) STRING enrichment analysis shows a protein– protein interaction network (21 nodes and 67 edges) of all enriched RNA-binding proteins and their associated biological processes.

We use the Transite set motif analysis tool to identify RNA-binding proteins (RBPs) recognized by the 3′ and 5′ untranslated regions (UTRs) of the common DEGs, which would indicate potential regulation of mRNA synthesis, translocation, and clearance. Three motifs within 3′ UTRs and 23 motifs within 5′ UTRs were significantly enriched in transcripts regulated by FAK-p130Cas signaling (**Fig. 4D**). Additionally, we used Cytoscape software to construct a protein-protein interaction (PPI) network with all enriched RBPs, revealing in a network composed of 21 nodes and 67 edges. A STRING (Search Tool for the Retrieval of Interacting Genes/Proteins) enrichment analysis of the PPI network indicated that the node RBPs were associated with the 3′ and 5′ UTRs (see **Fig. 4D**) and all linked to biological processes highlighted in the network, including “regulation of RNA metabolic process,” “RNA splicing,” “regulation of translation,” “macromolecule metabolic process,” and “regulation of cellular metabolic process” (**Fig. 4E**).

### FAK-p130Cas signaling affects lipid metabolic processes

To gain insight into the molecular consequences of interfering with FAK-p130Cas signaling, we used the Core Analysis function in the IPA software to analyze the differential expression of DEGs commonly regulated with FAK or p130Cas knockdown. There were differential and significant changes in six canonical pathways: five pathways were positively enriched and one pathway was negatively enriched (**Fig. 5A**). Pathways associated with “superpathway of cholesterol biosynthesis” and “PPAR signaling,” both associated with lipid metabolism, were positively and negatively enriched, respectively. The IPA Diseases and Functions feature identified 70 diseases and functions that were significantly and differentially impacted by FAK and p130Cas knockdown (**Fig. 5B**). Lipid metabolic processes, such as “lipolysis” (z-score, 3.249) and “metabolism of membrane lipid derivatives” (z-score, 2.734), were predicted to be highly activated. We then applied IPA’s Path Explorer tool to illustrate the molecular relationships among DEGs associated with lipolysis and metabolism of membrane lipid derivatives (**Fig. 5C, D**). The IPA Interaction Network Analysis identified three regulatory networks linking signaling events to metabolism: one associated with carbohydrate metabolism, nucleic acid metabolism, and small molecule biochemistry (p-score = 25) (**Fig. 5E**), one associated with lipid metabolism, small molecule biochemistry, and vitamin and mineral metabolism (p-score = 21) (**Fig. 5F**), and one associated with amino acid metabolism, cellular movement, and small molecule biochemistry (p-score = 21) (**Fig. 5G**). Collectively, these data demonstrate that inhibition of FAK-p130Cas signaling not only impacts lipid metabolism but also has far-reaching consequences on multiple metabolic pathways.

**Figure 5.**
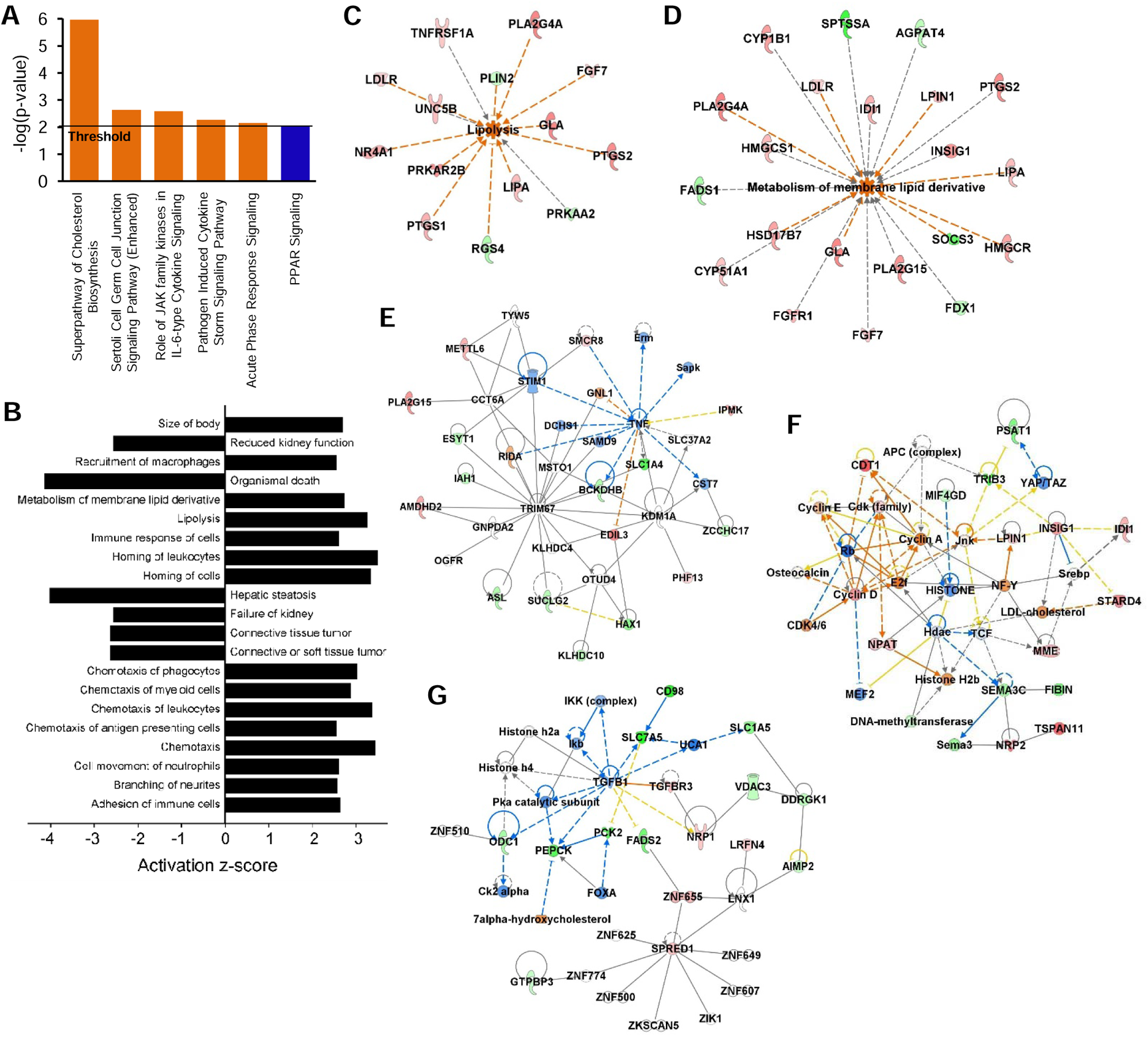
Inhibiting FAK-p130Cas signaling affects lipid metabolic processes. Histograms display differentially and significantly changed [absolute activation z score ≥ 2; -log_10_(*p* value) ≥ 2] canonical pathways (**A**) and diseases and functions (**B**) analyzed in QIAGEN IPA software using the Core Analysis function. Network diagrams acquired by IPA’s Path Explorer tool show relationships between 14 DEGs and lipolysis function (activation z-score, 3.249) (**C**) and between 21 DEGs and metabolism of membrane lipid derivate (activation z-score, 2.734) (**D**). IPA Interaction Network Analysis identified three regulatory networks associated with carbohydrate metabolism, nucleic acid metabolism, and small molecule biochemistry (p-score, 25) (**E**), lipid metabolism, small molecule biochemistry, and vitamin and mineral metabolism (p-score, 21) (**F**), and amino acid metabolism, cellular movement, and small molecule biochemistry (p-score, 21) (**G**).

### FAK-p130Cas signaling affects various metabolic processes via hub genes

We constructed another PPI network to identify highly connected hub genes—genes that have numerous interactions with other genes—among commonly regulated DEGs in response to both FAK and p130Cas knockdown. The resultant PPI network comprised 213 nodes and 649 edges (**Fig. S3**). We then performed k-means clustering by using the STRING online tool, which identified four functionally distinct clusters within the PPI network (**Fig. 6A**). Cluster 1, with 28 nodes and 96 edges (**Fig. 6B**), was linked to “amino acid metabolism,” “alpha-amino acid metabolism,” “alpha-amino acid biosynthetic process,” “carboxylic acid metabolic process,” etc. (**Fig. 6C**). Cluster 2, comprising 54 nodes and 55 edges (**Fig. 6D**), was enriched in “anatomical structure of morphogenesis,” “regulation of cell migration,” “cell migration,” “blood vessel development,” etc. (**Fig. 6E**). Cluster 3, with 33 nodes and 61 edges (**Fig. 6F**), was associated with “lipid biosynthetic process,” “sterol metabolic process,” “lipid metabolic process,” “sterol biosynthetic process,” etc. (**Fig. 6G**). Cluster 4, comprising 73 nodes and 264 edges (**Fig. 6H**), was enriched in “negative regulation of cellular process,” “anatomical structure development,” “positive regulation of biological process,” “regulation of protein metabolic process,” etc. (**Fig. 6I**).

**Figure 6.**
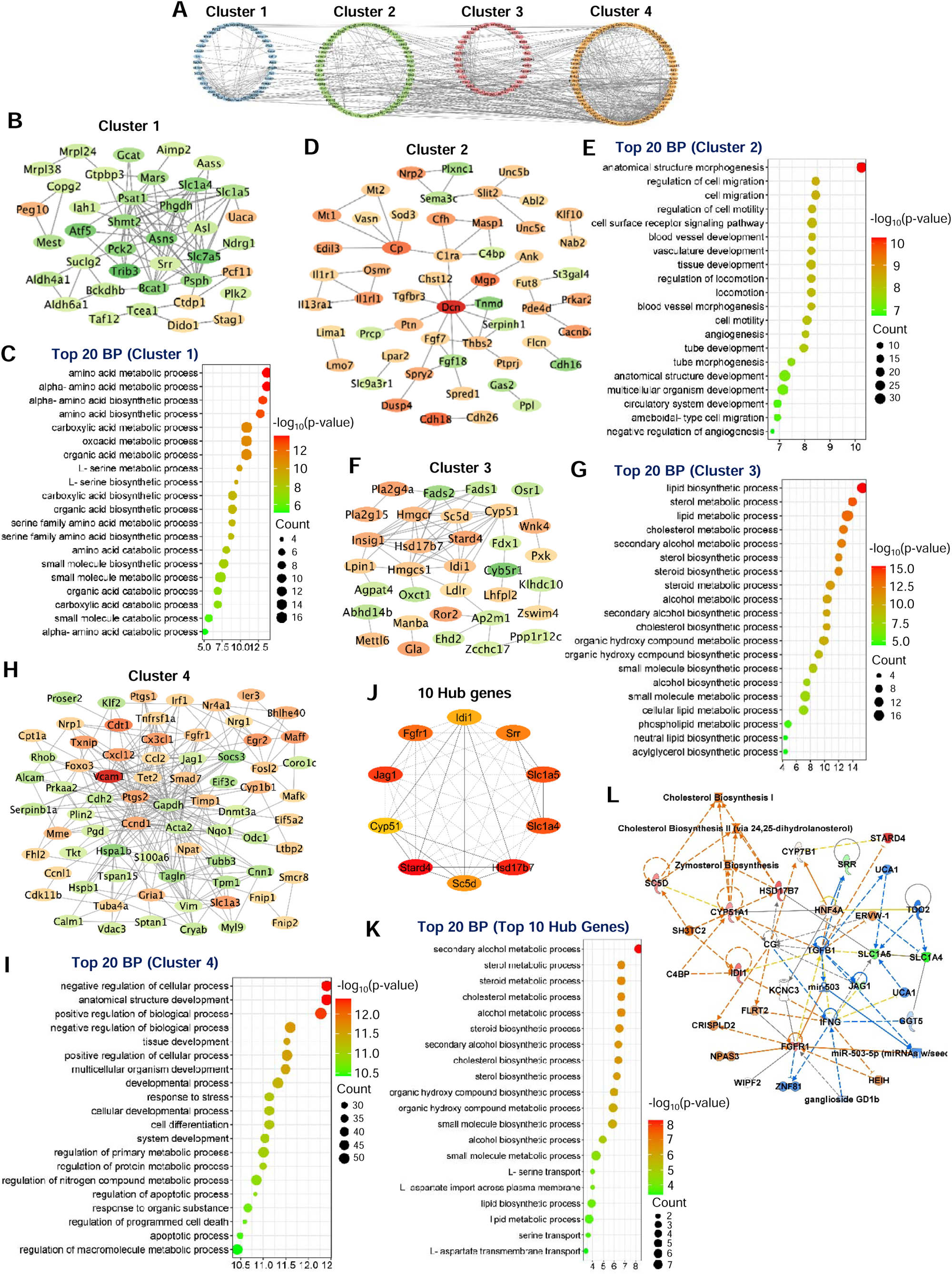
Hub genes mediate the impact of FAK and p130Cas knockdown on various metabolic processes. (**A**) Four clusters within the protein–protein interaction network of commonly regulated genes in FAK-p130Cas signaling. (**B-I**) The protein–protein interaction networks of the four identified clusters (**B, D, F, and H**) and bubble plots (**C, E, G, and I**) illustrate the top 20 enriched biological processes corresponding to each network. (**J**) Network (10 nodes and 45 edges) displays the top 10 hub genes identified using the Density of Maximum Neighborhood Component topological algorithm in the cytoHubba plug-in. (**K**) Bubble plot shows top 20 enriched biological processes of the top 10 hub genes. (**L**) The hub gene network (p-score, 32) associates with lipid metabolism, small molecule biosynthesis, and vitamin and mineral metabolism.

We identified the top ten hub genes within the PPI network for DEGs commonly regulated by both FAK and p130Cas knockdown in comparison to expression in control siRNA-treated cells on stiff hydrogels by using the Density of Maximum Neighborhood Component (DMNC) topological algorithm of the cytoHubba application. This analysis identified *Hsd17b7*, *Slc1a4*, *Srr*, *Idi1*, *Stard4*, *Jag1*, *Cyp51*, *Slc1a5*, *Fgfr1*, and *Sc5d* as highly connected hub genes within the PPI network (**Fig. 6J**). These ten hub genes are involved in various metabolic and biosynthetic processes, including “secondary alcohol metabolic process,” “steroid metabolic process,” “cholesterol metabolic process,” “cholesterol biosynthetic processes,” and “small molecule biosynthetic process” (**Fig. 6K**). An IPA core analysis of the expression log ratios of these ten genes revealed a hub gene network (p-score, 32) associated with lipid metabolism, small molecule biochemistry, and vitamin and mineral metabolism (**Fig. 6L**). Thus, the hub genes, intricately connected within the network, play central roles in various metabolic processes affected by inhibition of FAK-p130Cas signaling.

## DISCUSSION

In this study, we investigated the impact of focal adhesion proteins (FAK and p130Cas) on the early transcriptional responses of MEFs to ECM stiffness. These studies support and extend the results of previous studies, which showed that adhesion-mediated mechanotransduction alters mitochondrial reprogramming [9] as well as actin cytoskeletal dynamics [40, 41] and sterol responsive element binding protein (SREBP) activity [42]. Our findings reveal that ECM stiffness triggers robust transcriptional changes associated with metabolic, biosynthetic, and developmental processes within 1 h, indicating a mechanism for rapid bioenergetic changes in MEFs in response to changes in substrate stiffness.

FAK is activated by ECM/integrin interactions, and ECM stiffness induces metastasis and autophagy of cancer cells through integrin-FAK signaling [43, 44]. FAK activity in solid tumors leads to an increase in intracellular glucose levels as a result of a shift from OXPHOS (mitochondrial oxidative phosphorylation) to glycolysis [45]. Our findings show that in normal cells, FAK depletion (via siRNA knockdown) significantly impacts metabolic, biosynthetic, and developmental cell processes. FAK depletion in cancer-associated fibroblasts triggers an upregulation of CC chemokines (or β-chemokines), resulting in an increase in malignant cell glycolysis [46]. The KEGG pathway analysis in the present study indicates that the changes with FAK depletion are linked to metabolic disorders (diabetic cardiomyopathy), cancers (pancreatic, breast, lung, etc.), and neurodegenerative diseases (Alzheimer’s, Parkinson’s, and Huntington diseases), all involving altered metabolism and biosynthesis [47–53]. Altogether, the results from our study highlight the importance of FAK in regulating crucial metabolic and biosynthetic processes.

Cancer cells use signaling pathways, such as those involving ERK1/2, MAPK, Akt, and Rap1, to switch to aerobic glycolysis (known as the Warburg effect) to facilitate rapid proliferation [54]. Rap1 deficiency in mice alters the expression of genes involved in metabolism, leading to obesity and glucose intolerance [55]. These same signaling molecules are regulated by p130Cas [56–59], and previous studies as well as the data we present here indicate that p130Cas is crucial for modulating intracellular responses in response to changes in ECM stiffness [21, 60]. Specifically, our studies reveal a previously unrecognized role of p130Cas as a regulator of transcriptional programs involving metabolic and biosynthetic processes, such that information about ECM stiffness is mechanotransduced via p130Cas signaling to modulate transcriptional programs for various metabolic and biosynthetic processes.

The interaction between FAK and p130Cas is key for the early transcriptional responses of cells to their microenvironment. Inhibition of the FAK-p130Cas signaling resulted in an upregulation of *Il1r1* (interleukin 1 receptor, type I) and *Tnfrsf1a* (tumor necrosis factor receptor superfamily member 1A; also known as TNFR1), which reduce peroxisome-proliferator-activated receptor (PPAR) signaling [61]. PPAR signaling regulates the expression of enzymes involved in intracellular lipid metabolic processes, such as fatty acid uptake, lipid homeostasis, and peroxisome proliferation [62, 63]. Interestingly, we found that knockdown-mediated inhibition of FAK-p130Cas signaling upregulated genes for cyclooxygenase enzymes (*Ptgs1* and *Ptgs2*), nine other genes necessary for lipolysis [64–73], and genes enriched in various metabolic processes. The data indicate that stiffness-mediated early transcriptional responses in cells with knockdown of FAK-p130Cas signaling alter lipid metabolism by impairing PPAR signaling and fatty acid utilization. This leads cells to activate the superpathway of cholesterol metabolism and biosynthesis signaling, producing more cholesterol in place of the unused fatty acids.

We found that untranslated regions of common transcripts regulated by FAK and p130Cas are enriched for RBP motifs. For example, the 3′ UTR motifs are recognized by PTBP1, PTBP2, ROD1, SFRS1, and ZRANB2, which are important for stabilizing mRNAs by promoting exon inclusion [74] and repressing premature slicing and degradation [75]. Additionally, the common 5′ motifs recognized AU-rich elements such as ELAVL proteins, which stabilize mRNAs; e.g., ELAVL1 stabilizes *Ptgs2* mRNA in colon cancer cell lines, which causes an increase in COX-2 levels [76]. The negative enrichment for 5′ UTR motifs recognizing TIA1 and ZC3H10, which promote mRNA storage and degradation [77, 78], is further evidence that the transcriptional response mediated by FAK-p130Cas signaling involves translational regulation; however, further studies are needed to assess actual translation efficiency.

Our analyses with the cytoHubba application of Cytoscape software identified ten hub DEGs common to FAK and p130Cas knockdown. These genes, *Hsd17b7*, *Slc1a4*, *Srr*, *Idi1*, *Stard4*, *Jag1*, *Cyp51*, *Slc1a5*, *Fgfr1*, and *Sc5d*, are mainly associated with metabolic and biosynthetic processes. The downregulation of *Slc1a4* and *Slc1a5*, which code for glutamine transporters, with FAK-p130Cas knockdown is interesting, because cancer cells rely on the transport of glutamine through these receptors to meet their energy demands [79]. Many regulators of cholesterol metabolism were among the hub genes that were upregulated with FAK and p130Cas knockdown. These include the genes for Stard4, which is known to regulate intracellular cholesterol homeostasis [80], Hsd17b7, which is an enzyme that produces zymosterol [81], a precursor for cholesterol biosynthesis, and estradiol [82]. Sc5d catalyzes the synthesis of 7-dehydrocholesterol [83] to form cholesterol, and Idi1 and Cyp51 are part of the superpathway of the canonical cholesterol biosynthesis pathway [84].

## CONCLUSION

This study provides new insights into the genome-wide early transcriptional changes mediated by FAK-p130Cas signaling in response to ECM stiffness. FAK and p130Cas knockdown altered transcriptomes for metabolic reprogramming, increasing the expression of genes involved in cholesterol biosynthesis and reducing the expression of those involved in PPAR signaling and fatty acid utilization. Our analyses identified ten hub genes that were common targets of FAK and p130Cas signaling and are regulators of metabolic and biosynthetic processes. Altogether, the findings presented here reveal an important role of FAK and p130Cas as mechanotransduction signals to modulate metabolic processes enabling cells to adapt to the bioenergetic demands aroused by ECM stiffness.

## ACKNOWLEDGMENTS

We thank Karen Dietz for critical reading and editing of the manuscript. This work was supported by NIH/NHLBI grant (R01HL163168) to Y.B.

## AUTHOR CONTRIBUTIONS

Conceptualization, B.T., V.M.T., J.A.B., and Y.B.; methodology, B.T. V.M.T., and Y.B.; software, B.T.; investigation, B.T. and Y.B.; resources, Y.B.; writing – original draft, B.T. and Y.B.; writing – review & editing, B.T., V.M.T., J.A.B., P.Y., and Y.B.; visualization, B.T. and Y.B.; supervision, Y.B.; funding acquisition, Y.B.

## SUPPLEMENTARY INFORMATION

**Figure S1.**
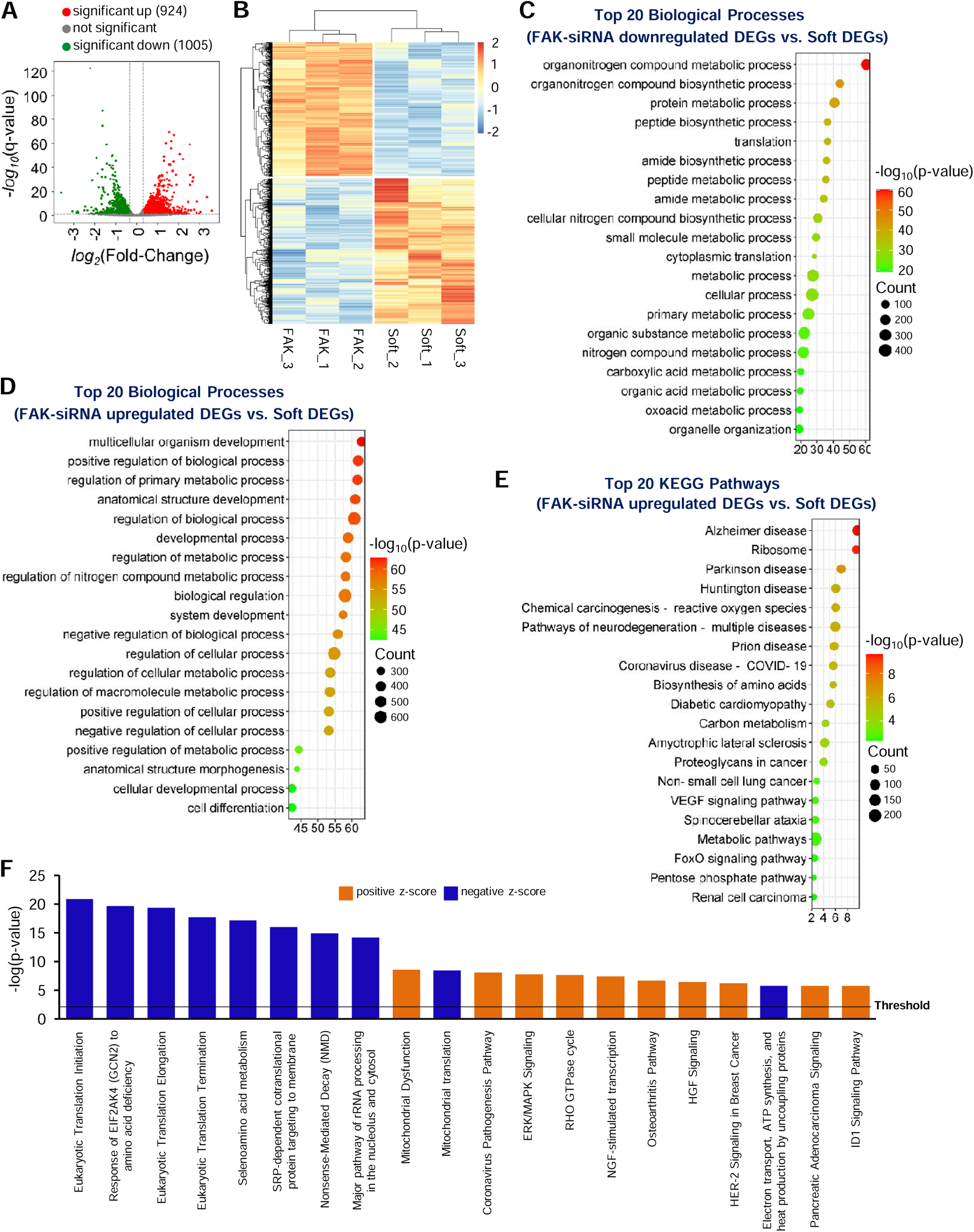
Transcriptional changes related to metabolism, biosynthesis, development, and signaling pathways in cells with FAK knockdown placed on soft hydrogels. Volcano plot (**A**) and heat map (**B**) to visualize the distribution and expression patterns of DEGs between FAK siRNA-transfected cells on stiff hydrogels and control siRNA-treated cells on soft hydrogels. Bubble plots show the top 20 enriched biological processes for significantly downregulated (**C**) and upregulated DEGs (**D**) comparing cells on stiff hydrogels with FAK siRNA to cells on soft hydrogels with control siRNA. (**E**) Top 20 enriched KEGG pathways among DEGs in cells on stiff hydrogels with FAK siRNA versus cells on soft hydrogels with control siRNA. (**F**) Histogram represents the top 20 significantly and differentially activated or inhibited [absolute activation z score ≥ 2; -log_10_(*p* value) ≥ 2] canonical pathways in response to FAK knockdown.

**Figure S2.**
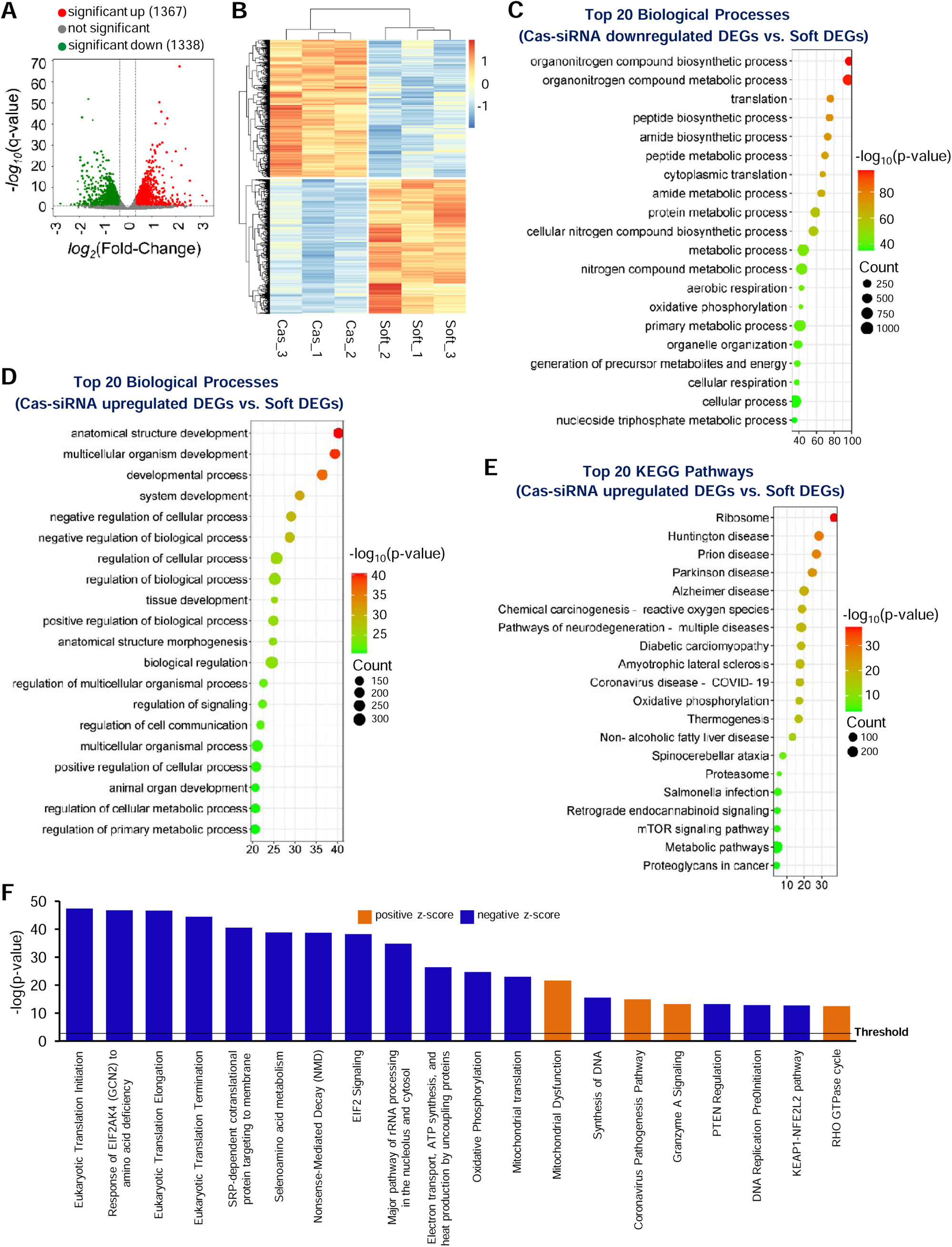
Transcriptional changes related to metabolism, biosynthesis, development, and disease in cells with p130Cas knockdown placed on soft hydrogels. Volcano plot (**A**) and heat map (**B**) to visualize the distribution and expression patterns of DEGs between p130Cas siRNA-transfected cells on stiff hydrogels and control siRNA-treated cells on soft hydrogels. Bubble plots show the top 20 enriched biological processes for significantly downregulated (**C**) and upregulated (**D**) DEGs comparing cells on stiff hydrogels with p130Cas siRNA to cells on soft hydrogels with control siRNA. (**E**) Top 20 enriched KEGG pathways among DEGs in cells on stiff hydrogels with p130Cas siRNA versus cells on soft hydrogels with control siRNA. (**F**) Histogram represents the top 20 significantly and differentially activated or inhibited [absolute activation z score ≥ 2; -log_10_(*p* value) ≥ 2] canonical pathways in response to p130Cas knockdown.

**Figure S3.**
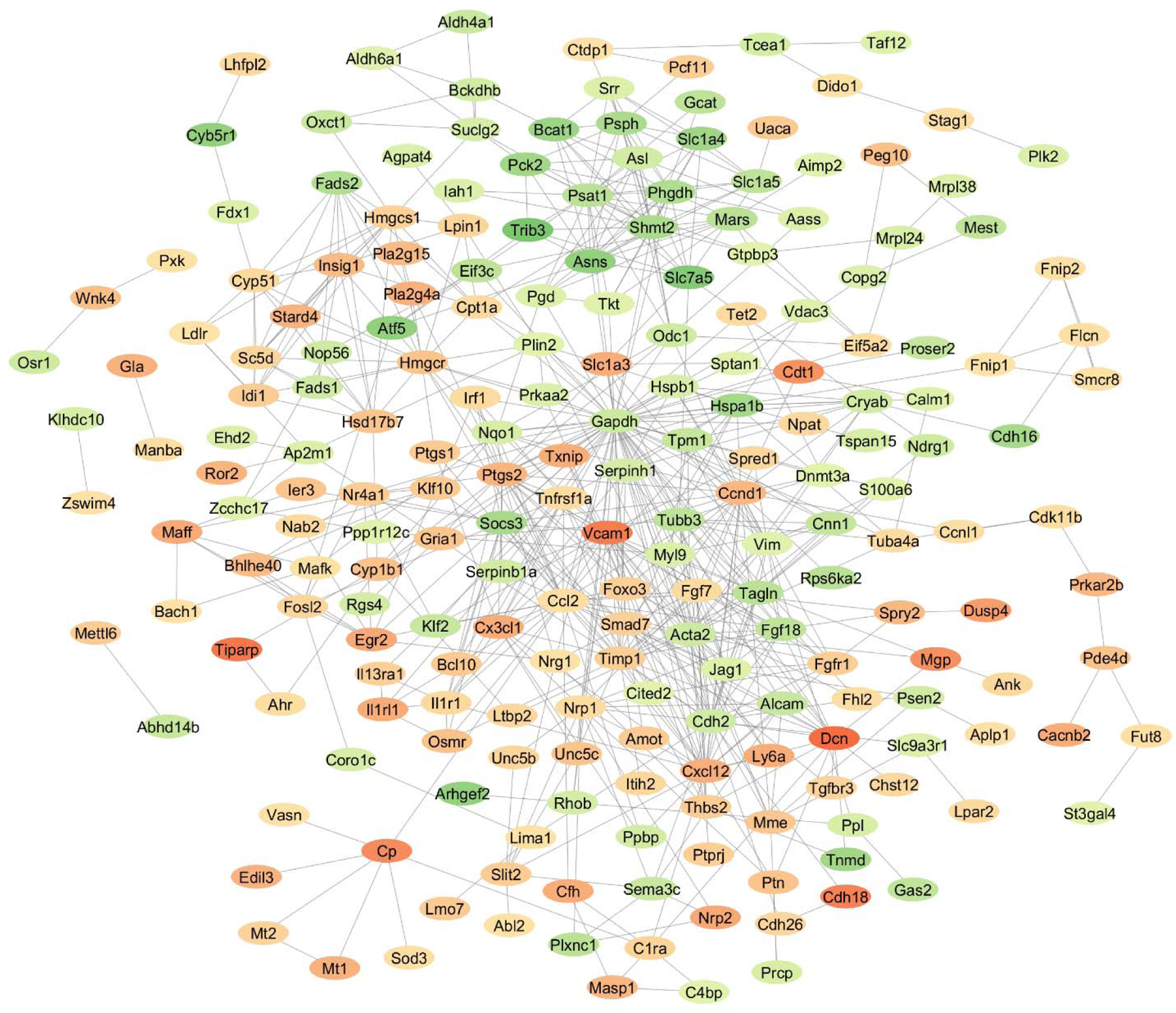
Protein–protein interaction network (213 nodes and 649 edges) of commonly regulated genes in FAK-p130Cas signaling.

## REFERENCES

1. Zanotelli, M.R., Rahman-Zaman, A., VanderBurgh, J.A. et al. Energetic costs regulated by cell mechanics and confinement are predictive of migration path during decision-making. Nat Commun 10, 4185 (2019). 10.1038/s41467-019-12155-z

2. Xie, Jing et al. “Energy expenditure during cell spreading influences the cellular response to matrix stiffness.” Biomaterials vol. 267 (2021): 120494. doi:10.1016/j.biomaterials.2020.120494

3. Irène Dang, Joseph A. Brazzo, Yongho Bae, Richard K. Assoian; Key role for Rac in the early transcriptional response to ECM stiffness and the stiffness-dependent repression of ATF3. J Cell Sci 2023; jcs.260636. doi: 10.1242/jcs.260636 Bartolák-Suki, E., Imsirovic, J., Parameswaran, H. et al. Fluctuation-driven mechanotransduction regulates mitochondrial-network structure and function. Nature Mater 14, 1049–1057 (2015). 10.1038/nmat4358

4. Biber, John C et al. “Survivin as a mediator of stiffness-induced cell cycle progression and proliferation of vascular smooth muscle cells.” APL bioengineering vol. 7,4 046108. 30 Oct. 2023, doi:10.1063/5.0150532

5. Sun, Meiyu et al. “Extracellular matrix stiffness controls osteogenic differentiation of mesenchymal stem cells mediated by integrin α5.” Stem cell research & therapy vol. 9,1 52. 1 Mar. 2018, doi:10.1186/s13287-018-0798-0

6. Discher, Dennis E et al. “Tissue cells feel and respond to the stiffness of their substrate.” Science (New York, N.Y.) vol. 310,5751 (2005): 1139–43. doi:10.1126/science.1116995

7. Ge H, Tian M, Pei Q, Tan F, Pei H. Extracellular Matrix Stiffness: New Areas Affecting Cell Metabolism. Front Oncol. 2021 Feb 24;11:631991. doi: 10.3389/fonc.2021.631991. PMID: 33718214; PMCID: PMC7943852.

8. Taufalele, Paul V, and Cynthia A Reinhart-King. “Matrix stiffness primes cells for future oxidative stress.” Trends in cancer vol. 7,10 (2021): 883–885. doi:10.1016/j.trecan.2021.08.003

9. Tharp, Kevin M et al. “Adhesion-mediated mechanosignaling forces mitohormesis.” Cell metabolism vol. 33,7 (2021): 1322–1341.e13. doi:10.1016/j.cmet.2021.04.017

10. Ata, Rehman, and Costin N Antonescu. “Integrins and Cell Metabolism: An Intimate Relationship Impacting Cancer.” International journal of molecular sciences vol. 18,1 189. 18 Jan. 2017, doi:10.3390/ijms18010189

11. Guan, J L. “Role of focal adhesion kinase in integrin signaling.” The international journal of biochemistry & cell biology vol. 29,8-9 (1997): 1085–96. doi:10.1016/s1357-2725(97)00051-4

12. Lawson C, Lim ST, Uryu S, Chen XL, Calderwood DA, Schlaepfer DD. FAK promotes recruitment of talin to nascent adhesions to control cell motility. J Cell Biol. 2012 Jan 23;196(2):223–32. doi: 10.1083/jcb.201108078. Erratum in: J Cell Biol. 2012 Feb 6;196(3):387. PMID: 22270917; PMCID: PMC3265949.

13. Schaller, M D et al. “Autophosphorylation of the focal adhesion kinase, pp125FAK, directs SH2-dependent binding of pp60src.” Molecular and cellular biology vol. 14,3 (1994): 1680–8. doi:10.1128/mcb.14.3.1680-1688.1994

14. Brazzo, Joseph A et al. “Mechanosensitive expression of lamellipodin promotes intracellular stiffness, cyclin expression and cell proliferation.” Journal of cell science vol. 134,12 (2021): jcs257709. doi:10.1242/jcs.257709

15. Defilippi, Paola et al. “p130Cas: a versatile scaffold in signaling networks.” Trends in cell biology vol. 16,5 (2006): 257–63. doi:10.1016/j.tcb.2006.03.003

16. Bae, Yong Ho et al. “A FAK-Cas-Rac-lamellipodin signaling module transduces extracellular matrix stiffness into mechanosensitive cell cycling.” Science signaling vol. 7,330 ra57. 17 Jun. 2014, doi:10.1126/scisignal.2004838

17. Pratt SJ, Epple H, Ward M, Feng Y, Braga VM, Longmore GD. The LIM protein Ajuba influences p130Cas localization and Rac1 activity during cell migration. J Cell Biol. 2005 Feb 28;168(5):813–24. doi: 10.1083/jcb.200406083. Epub 2005 Feb 22. PMID: 15728191; PMCID: PMC2171823.

18. Sharma, A., Mayer, B.J. Phosphorylation of p130Cas initiates Rac activation and membrane ruffling. BMC Cell Biol 9, 50 (2008). 10.1186/1471-2121-9-50

19. Hupfer, Anna et al. “Matrix stiffness drives stromal autophagy and promotes formation of a protumorigenic niche.” Proceedings of the National Academy of Sciences of the United States of America vol. 118,40 (2021): e2105367118. doi:10.1073/pnas.2105367118

20. Hsia DA, Mitra SK, Hauck CR, Streblow DN, Nelson JA, Ilic D, Huang S, Li E, Nemerow GR, Leng J, Spencer KS, Cheresh DA, Schlaepfer DD. Differential regulation of cell motility and invasion by FAK. J Cell Biol. 2003 Mar 3;160(5):753–67. doi: 10.1083/jcb.200212114. PMID: 12615911; PMCID: PMC2173366.

21. Cary, L A et al. “Identification of p130Cas as a mediator of focal adhesion kinase-promoted cell migration.” The Journal of cell biology vol. 140,1 (1998): 211–21. doi:10.1083/jcb.140.1.211

22. Mitra, Satyajit K, and David D Schlaepfer. “Integrin-regulated FAK-Src signaling in normal and cancer cells.” Current opinion in cell biology vol. 18,5 (2006): 516–23. doi:10.1016/j.ceb.2006.08.011

23. Cretu A, Castagnino P, Assoian R. Studying the effects of matrix stiffness on cellular function using acrylamide-based hydrogels. J Vis Exp. 2010;(42):2089. Published 2010 Aug 10. doi:10.3791/2089

24. Babraham Bioinformatics – FastQC A Quality Control tool for High Throughput Sequence Data. http://www.bioinformatics.babraham.ac.uk/projects/fastqc/ [August 21, 2018]

25. Wingett SW, Andrews S. FastQ Screen: A tool for multi-genome mapping and quality control. F1000Res. 2018 Aug 24;7:1338. doi: 10.12688/f1000research.15931.2. PMID: 30254741; PMCID: PMC6124377.

26. Ewels, Philip et al. “MultiQC: summarize analysis results for multiple tools and samples in a single report.” Bioinformatics (Oxford, England) vol. 32,19 (2016): 3047–8. doi:10.1093/bioinformatics/btw354

27. Kim, D., Langmead, B. & Salzberg, S. HISAT: a fast spliced aligner with low memory requirements. Nat Methods 12, 357–360 (2015). 10.1038/nmeth.3317

28. Liao, Yang et al. “featureCounts: an efficient general purpose program for assigning sequence reads to genomic features.” Bioinformatics (Oxford, England) vol. 30,7 (2014): 923–30. doi:10.1093/bioinformatics/btt656

29. Wickham H (2016). ggplot2: Elegant Graphics for Data Analysis. Springer-Verlag New York. ISBN 978-3-319-24277-4, https://ggplot2.tidyverse.org.

30. Kolde, Raivo. “Pheatmap: pretty heatmaps.” R package version 1.2 (2012): 726.

31. Love MI, Huber W, Anders S (2014). “Moderated estimation of fold change and dispersion for RNA-seq data with DESeq2.” Genome Biology, 15, 550. doi:10.1186/s13059-014-0550-8.

32. Renesh Bedre. (2020, March 5). reneshbedre/bioinfokit: Bioinformatics data analysis and visualization toolkit. Zenodo. 10.5281/zenodo.3698145.

33. Reimand J, Kull M, Peterson H, Hansen J, Vilo J. g:Profiler--a web-based toolset for functional profiling of gene lists from large-scale experiments. Nucleic Acids Res. 2007 Jul;35(Web Server issue):W193–200. doi: 10.1093/nar/gkm226. Epub 2007 May 3. PMID: 17478515; PMCID: PMC1933153.

34. Krämer, A., Green, J., Pollard, Jr., J., and Tugendreich, S. (2014) Causal analysis approaches in Ingenuity Pathway Analysis. Bioinformatics. 30(4):523–30

35. Bindea G, Mlecnik B, Hackl H, Charoentong P, Tosolini M, Kirilovsky A, Fridman WH, Pagès F, Trajanoski Z, Galon J. ClueGO: a Cytoscape plug-in to decipher functionally grouped gene ontology and pathway annotation networks. Bioinformatics. 2009 Apr 15;25(8):1091–3. doi: 10.1093/bioinformatics/btp101. Epub 2009 Feb 23. PMID: 19237447; PMCID: PMC2666812.

36. Szklarczyk, Damian et al. “STRING v10: protein-protein interaction networks, integrated over the tree of life.” Nucleic acids research vol. 43,Database issue (2015): D447–52. doi:10.1093/nar/gku1003

37. Chin, Chia-Hao et al. “cytoHubba: identifying hub objects and sub-networks from complex interactome.” BMC systems biology vol. 8 Suppl 4,Suppl 4 (2014): S11. doi:10.1186/1752-0509-8-S4-S11

38. Krismer, Konstantin et al. “Transite: A Computational Motif-Based Analysis Platform That Identifies RNA-Binding Proteins Modulating Changes in Gene Expression.” Cell reports vol. 32,8 (2020): 108064. doi:10.1016/j.celrep.2020.108064

39. Shannon P, Markiel A, Ozier O, et al. Cytoscape: a software environment for integrated models of biomolecular interaction networks. Genome Res. 2003;13(11):2498–2504. Doi:10.1101/gr.1239303

40. Liu, Zhuo et al. “The role of cytoskeleton in glucose regulation.” Biochemistry. Biokhimiia vol. 71,5 (2006): 476–80. doi:10.1134/s0006297906050026

41. Park, Jin Suk et al. “Mechanical regulation of glycolysis via cytoskeleton architecture.” Nature vol. 578,7796 (2020): 621–626. doi:10.1038/s41586-020-1998-1

42. Romani, Patrizia et al. “Extracellular matrix mechanical cues regulate lipid metabolism through Lipin-1 and SREBP.” Nature cell biology vol. 21,3 (2019): 338–347. doi:10.1038/s41556-018-0270-5

43. Murphy, J.M., Rodriguez, Y.A.R., Jeong, K. et al. Targeting focal adhesion kinase in cancer cells and the tumor microenvironment. Exp Mol Med 52, 877–886 (2020). 10.1038/s12276-020-0447-4

44. Chuang HH, Zhen YY, Tsai YC, Chuang CH, Hsiao M, Huang MS, Yang CJ. FAK in Cancer: From Mechanisms to Therapeutic Strategies. Int J Mol Sci. 2022 Feb 2;23(3):1726. doi: 10.3390/ijms23031726. PMID: 35163650; PMCID: PMC8836199

45. Zhang, J et al. “Focal adhesion kinase-promoted tumor glucose metabolism is associated with a shift of mitochondrial respiration to glycolysis.” Oncogene vol. 35,15 (2016): 1926–42. doi:10.1038/onc.2015.256

46. Demircioglu, Fevzi et al. “Cancer associated fibroblast FAK regulates malignant cell metabolism.” Nature communications vol. 11,1 1290. 10 Mar. 2020, doi:10.1038/s41467-020-15104-3

47. Chong CR, Clarke K, Levelt E. Metabolic Remodeling in Diabetic Cardiomyopathy. Cardiovasc Res. 2017;113(4):422–430. doi:10.1093/cvr/cvx018

48. Encarnación-Rosado J, Kimmelman AC. Harnessing metabolic dependencies in pancreatic cancers. Nat Rev Gastroenterol Hepatol. 2021;18(7):482–492. doi:10.1038/s41575-021-00431-7

49. Wang Z, Jiang Q, Dong C. Metabolic reprogramming in triple-negative breast cancer. Cancer Biol Med. 2020;17(1):44–59. doi:10.20892/j.issn.2095-3941.2019.0210

50. Vanhove K, Graulus GJ, Mesotten L, et al. The Metabolic Landscape of Lung Cancer: New Insights in a Disturbed Glucose Metabolism. Front Oncol. 2019;9:1215. Published 2019 Nov 15. doi:10.3389/fonc.2019.01215

51. Kang S, Lee YH, Lee JE. Metabolism-Centric Overview of the Pathogenesis of Alzheimer’s Disease. Yonsei Med J. 2017;58(3):479–488. doi:10.3349/ymj.2017.58.3.479

52. Sonninen TM, Hämäläinen RH, Koskuvi M, et al. Metabolic alterations in Parkinson’s disease astrocytes. Sci Rep. 2020;10(1):14474. Published 2020 Sep 2. doi:10.1038/s41598-020-71329-8

53. Nambron R, Silajdžić E, Kalliolia E, et al. A Metabolic Study of Huntington’s Disease. PLoS One. 2016;11(1):e0146480. Published 2016 Jan 8. doi:10.1371/journal.pone.0146480

54. Papa S, Choy PM, Bubici C. The ERK and JNK pathways in the regulation of metabolic reprogramming. Oncogene. 2019;38(13):2223–2240. doi:10.1038/s41388-018-0582-8

55. Yeung F, Ramírez CM, Mateos-Gomez PA, et al. Nontelomeric role for Rap1 in regulating metabolism and protecting against obesity. Cell Rep. 2013;3(6):1847–1856. doi:10.1016/j.celrep.2013.05.032

56. Cabodi S, Moro L, Baj G, et al. p130Cas interacts with estrogen receptor alpha and modulates non-genomic estrogen signaling in breast cancer cells. J Cell Sci. 2004;117(Pt 8):1603–1611. Doi:10.1242/jcs.01025

57. Kodama H, Fukuda K, Takahashi E, et al. Selective involvement of p130Cas/Crk/Pyk2/c-Src in endothelin-1-induced JNK activation. Hypertension. 2003;41(6):1372–1379. doi:10.1161/01.HYP.0000069698.11814.F4

58. Costamagna A, Natalini D, Camacho Leal MDP, et al. Docking Protein p130Cas Regulates Acinar to Ductal Metaplasia During Pancreatic Adenocarcinoma Development and Pancreatitis. Gastroenterology. 2022;162(4):1242–1255.e11. doi:10.1053/j.gastro.2021.12.242

59. Gotoh T, Cai D, Tian X, Feig LA, Lerner A. p130Cas regulates the activity of AND-34, a novel Ral, Rap1, and R-Ras guanine nucleotide exchange factor. J Biol Chem. 2000;275(39):30118–30123. doi:10.1074/jbc.M003074200

60. Sawada Y, Tamada M, Dubin-Thaler BJ, et al. Force sensing by mechanical extension of the Src family kinase substrate p130Cas. Cell. 2006;127(5):1015–1026. doi:10.1016/j.cell.2006.09.044

61. Kim MS, Sweeney TR, Shigenaga JK, Chui LG, Moser A, Grunfeld C, Feingold KR. Tumor necrosis factor and interleukin 1 decrease RXRalpha, PPARalpha, PPARgamma, LXRalpha, and the coactivators SRC-1, PGC-1alpha, and PGC-1beta in liver cells. Metabolism. 2007 Feb;56(2):267–79. doi: 10.1016/j.metabol.2006.10.007. PMID: 17224343; PMCID: PMC2700944.

62. Grygiel-Górniak, Bogna. “Peroxisome proliferator-activated receptors and their ligands: nutritional and clinical implications--a review.” Nutrition journal vol. 13 17. 14 Feb. 2014, doi:10.1186/1475-2891-13-17

63. Gervois, P et al. “Regulation of lipid and lipoprotein metabolism by PPAR activators.” Clinical chemistry and laboratory medicine vol. 38,1 (2000): 3–11. doi:10.1515/CCLM.2000.002

64. Smith, W L et al. “Cyclooxygenases: structural, cellular, and molecular biology.” Annual review of biochemistry vol. 69 (2000): 145–82. doi:10.1146/annurev.biochem.69.1.145

65. Carter, C J. “Convergence of genes implicated in Alzheimer’s disease on the cerebral cholesterol shuttle: APP, cholesterol, lipoproteins, and atherosclerosis.” Neurochemistry international vol. 50,1 (2007): 12–38. doi:10.1016/j.neuint.2006.07.007

66. Planas, J V et al. “Mutation of the RIIbeta subunit of protein kinase A differentially affects lipolysis but not gene induction in white adipose tissue.” The Journal of biological chemistry vol. 274,51 (1999): 36281–7. doi:10.1074/jbc.274.51.36281

67. Maxwell, Megan A et al. “Nur77 regulates lipolysis in skeletal muscle cells. Evidence for cross-talk between the beta-adrenergic and an orphan nuclear hormone receptor pathway.” The Journal of biological chemistry vol. 280,13 (2005): 12573–84. doi:10.1074/jbc.M409580200

68. Nonogaki, K et al. “Keratinocyte growth factor increases fatty acid mobilization and hepatic triglyceride secretion in rats.” Endocrinology vol. 136,10 (1995): 4278–84. doi:10.1210/endo.136.10.7664645

69. Xu, You-Hai et al. “Multi-system disorders of glycosphingolipid and ganglioside metabolism.” Journal of lipid research vol. 51,7 (2010): 1643–75. doi:10.1194/jlr.R003996

70. Herz, J et al. “Initial hepatic removal of chylomicron remnants is unaffected but endocytosis is delayed in mice lacking the low density lipoprotein receptor.” Proceedings of the National Academy of Sciences of the United States of America vol. 92,10 (1995): 4611–5. doi:10.1073/pnas.92.10.4611

71. Bayburt, T, and M H Gelb. “Interfacial catalysis by human 85 kDa cytosolic phospholipase A2 on anionic vesicles in the scooting mode.” Biochemistry vol. 36,11 (1997): 3216–31. doi:10.1021/bi961659d

72. Geilen, C C et al. “1alpha,25-dihydroxyvitamin D3 induces sphingomyelin hydrolysis in HaCaT cells via tumor necrosis factor alpha.” The Journal of biological chemistry vol. 272,14 (1997): 8997–9001. doi:10.1074/jbc.272.14.8997

73. Xie, Yi et al. “DCC-dependent phospholipase C signaling in netrin-1-induced neurite elongation.” The Journal of biological chemistry vol. 281,5 (2006): 2605–11. doi:10.1074/jbc.M512767200

74. Paz S, Ritchie A, Mauer C, Caputi M. The RNA binding protein SRSF1 is a master switch of gene expression and regulation in the immune system. Cytokine Growth Factor Rev. 2021 Feb;57:19–26. doi: 10.1016/j.cytogfr.2020.10.008. Epub 2020 Nov 2. PMID: 33160830; PMCID: PMC7897272.

75. Licatalosi DD, Yano M, Fak JJ, Mele A, Grabinski SE, Zhang C, Darnell RB. Ptbp2 represses adult-specific splicing to regulate the generation of neuronal precursors in the embryonic brain. Genes Dev. 2012 Jul 15;26(14):1626–42. doi: 10.1101/gad.191338.112. PMID: 22802532; PMCID: PMC3404389.

76. Dixon DA, Tolley ND, King PH, Nabors LB, McIntyre TM, Zimmerman GA, Prescott SM. Altered expression of the mRNA stability factor HuR promotes cyclooxygenase-2 expression in colon cancer cells. J Clin Invest. 2001 Dec;108(11):1657–65. doi: 10.1172/JCI12973. PMID: 11733561; PMCID: PMC200983.

77. Díaz-Muñoz MD, Kiselev VY, Le Novère N, Curk T, Ule J, Turner M. Tia1 dependent regulation of mRNA subcellular location and translation controls p53 expression in B cells. Nat Commun. 2017 Sep 13;8(1):530. doi: 10.1038/s41467-017-00454-2. PMID: 28904350; PMCID: PMC5597594.

78. Audano M, Pedretti S, Ligorio S, Gualdrini F, Polletti S, Russo M, Ghisletti S, Bean C, Crestani M, Caruso D, De Fabiani E, Mitro N. Zc3h10 regulates adipogenesis by controlling translation and F-actin/mitochondria interaction. J Cell Biol. 2021 Mar 1;220(3):e202003173. doi: 10.1083/jcb.202003173. PMID: 33566069; PMCID: PMC7879490.

79. White MA, Lin C, Rajapakshe K, Dong J, Shi Y, Tsouko E, Mukhopadhyay R, Jasso D, Dawood W, Coarfa C, Frigo DE. Glutamine Transporters Are Targets of Multiple Oncogenic Signaling Pathways in Prostate Cancer. Mol Cancer Res. 2017 Aug;15(8):1017–1028. doi: 10.1158/1541-7786.MCR-16-0480. Epub 2017 May 15. PMID: 28507054; PMCID: PMC5685160.

80. Mesmin B, Pipalia NH, Lund FW, Ramlall TF, Sokolov A, Eliezer D, Maxfield FR. STARD4 abundance regulates sterol transport and sensing. Mol Biol Cell. 2011 Nov;22(21):4004–15. doi: 10.1091/mbc.E11-04-0372. Epub 2011 Sep 7. PMID: 21900492; PMCID: PMC3204063.

81. Pleshinger MJ, Friedrich RM, Hubler Z, Rivera-León AM, Gao F, Yan D, Sax JL, Srinivasan R, Bederman I, Shick HE, Tesar PJ, Adams DJ. Inhibition of SC4MOL and HSD17B7 shifts cellular sterol composition and promotes oligodendrocyte formation. RSC Chem Biol. 2021 Oct 21;3(1):56–68. doi: 10.1039/d1cb00145k. PMID: 35128409; PMCID: PMC8729178.

82. Nokelainen, P et al. “Expression cloning of a novel estrogenic mouse 17 beta-hydroxysteroid dehydrogenase/17-ketosteroid reductase (m17HSD7), previously described as a prolactin receptor-associated protein (PRAP) in rat.” Molecular endocrinology (Baltimore, Md.) vol. 12,7 (1998): 1048–59. doi:10.1210/mend.12.7.0134

83. Platt FM, Wassif C, Colaco A, Dardis A, Lloyd-Evans E, Bembi B, Porter FD. Disorders of cholesterol metabolism and their unanticipated convergent mechanisms of disease. Annu Rev Genomics Hum Genet. 2014;15:173–94. doi: 10.1146/annurev-genom-091212-153412. PMID: 25184529; PMCID: PMC6292211.

84. Rondini EA, Duniec-Dmuchowski Z, Cukovic D, Dombkowski AA, Kocarek TA. Differential Regulation of Gene Expression by Cholesterol Biosynthesis Inhibitors That Reduce (Pravastatin) or Enhance (Squalestatin 1) Nonsterol Isoprenoid Levels in Primary Cultured Mouse and Rat Hepatocytes. J Pharmacol Exp Ther. 2016 Aug;358(2):216–29. doi: 10.1124/jpet.116.233312. Epub 2016 May 25. PMID: 27225895; PMCID: PMC4959097.

