## Supplemental Tables for "FAK and p130Cas modulate stiffness-mediated early transcription and cellular metabolism"

**Table S1. RNA Sequencing Quality Control Analysis. ***

| | Initial QC | | | Analysis QC | | |
|---|---|---|---|---|---|---|
| Sample Name | % Dups | % GC | M Seqs | % Assigned | M Assigned | % Aligned |
| Cas2-High-Set1_pe_R1_001 | 83.30% | 60% | 57.1 | 31.90% | 19.8 | |
| Cas2-High-Set1_pe_R2_001 | 82.80% | 60% | 57.1 | | | 62.70% |
| Cas2-High-Set2_pe_R1_001 | 76.70% | 52% | 51.3 | 40.10% | 22.7 | |
| Cas2-High-Set2_pe_R2_001 | 74.40% | 52% | 51.3 | | | 82.10% |
| Cas2-High-Set3_pe_R1_001 | 80.40% | 55% | 60.3 | 46.70% | 30.7 | |
| Cas2-High-Set3_pe_R2_001 | 78.90% | 55% | 60.3 | | | 77.30% |
| Ctrl-High-Set1_pe_R1_001 | 81.00% | 53% | 58.6 | 41.30% | 26.9 | |
| Ctrl-High-Set1_pe_R2_001 | 78.60% | 53% | 58.6 | | | 81.50% |
| Ctrl-High-Set2_pe_R1_001 | 82.20% | 56% | 51.3 | 37.10% | 21.1 | |
| Ctrl-High-Set2_pe_R2_001 | 81.50% | 56% | 51.3 | | | 73.90% |
| Ctrl-High-Set3_pe_R1_001 | 77.90% | 53% | 59.7 | 45.00% | 29.4 | |
| Ctrl-High-Set3_pe_R2_001 | 77.00% | 53% | 59.7 | | | 82.90% |
| Ctrl-Low-Set1_pe_R1_001 | 82.10% | 58% | 43.7 | 36.10% | 17 | |
| Ctrl-Low-Set1_pe_R2_001 | 75.10% | 58% | 43.7 | | | 67.50% |
| Ctrl-Low-Set2_pe_R1_001 | 85.60% | 58% | 59.3 | 31.80% | 20.5 | |
| Ctrl-Low-Set2_pe_R2_001 | 84.90% | 58% | 59.3 | | | 66.40% |
| Ctrl-Low-Set3_pe_R1_001 | 82.20% | 55% | 58.1 | 42.90% | 27.3 | |
| Ctrl-Low-Set3_pe_R2_001 | 80.70% | 55% | 58.1 | | | 76.10% |
| FAK1-High-Set1_pe_R1_001 | 80.00% | 55% | 59.5 | 41.00% | 26.7 | |
| FAK1-High-Set1_pe_R2_001 | 79.30% | 56% | 59.5 | | | 75.70% |
| FAK1-High-Set2_pe_R1_001 | 81.40% | 55% | 65.9 | 44.20% | 31.7 | |
| FAK1-High-Set2_pe_R2_001 | 73.00% | 55% | 65.9 | | | 77.70% |
| FAK1-High-Set3_pe_R1_001 | 80.50% | 56% | 51.2 | 41.30% | 23.1 | |
| FAK1-High-Set3_pe_R2_001 | 79.80% | 56% | 51.2 | | | 74.60% |

* Quality of the RNA sequencing experiments. The sequencing experiments had an average of 56.3 million sequences per sample with an average of 74.9% read mapping rate. (Dups. = Duplicated, M. = Million, Seqs. = Sequences)

**Table S2. Gene Enrichment Results of Commonly Regulated DEGs by FAK and p130Cas knockdown**

| ID | Term | Term PValue BH-Corrected | Group PValue BH-Corrected | GOGroups | % Associated Genes | Nr. Genes |
|---|---|---|---|---|---|---|
| GO:0009987 | cellular process | 1.4716E-06 | 1.54061E-06 | Group00 | 1.135029316 | 203 |
| GO:0044281 | small molecule metabolic process | 6.6141E-13 | 4.34047E-13 | Group01 | 2.939632654 | 56 |
| GO:0006950 | response to stress | 5.1604E-11 | 9.76241E-15 | Group02 | 2.011225462 | 86 |
| GO:0009605 | response to external stimulus | 3.6384E-13 | 9.76241E-15 | Group02 | 2.388337374 | 77 |
| GO:0003008 | system process | 0.0053722 | 0.000109223 | Group03 | 1.4819206 | 50 |
| GO:0065008 | regulation of biological quality | 4.5759E-10 | 0.000109223 | Group03 | 1.986915469 | 82 |
| GO:0008283 | cell population proliferation | 8.8262E-09 | 7.90684E-09 | Group04 | 2.358700514 | 53 |
| GO:0042127 | regulation of cell population proliferation | 1.4141E-10 | 7.90684E-09 | Group04 | 2.711323738 | 51 |
| GO:0051179 | localization | 0.00754111 | 0.006463811 | Group05 | 1.323685169 | 75 |
| GO:0051234 | establishment of localization | 0.01077228 | 0.006463811 | Group05 | 1.335193276 | 67 |
| GO:0006810 | transport | 0.0189713 | 0.006463811 | Group05 | 1.318500161 | 64 |
| GO:0071840 | cellular component organization or biogenesis | 8.0453E-05 | 7.77106E-05 | Group06 | 1.423748732 | 97 |
| GO:0016043 | cellular component organization | 3.799E-05 | 7.77106E-05 | Group06 | 1.450151086 | 96 |
| GO:0051128 | regulation of cellular component organization | 3.9043E-06 | 7.77106E-05 | Group06 | 1.962293148 | 51 |
| GO:0006996 | organelle organization | 0.03676558 | 7.77106E-05 | Group06 | 1.314810276 | 53 |
| GO:0040011 | locomotion | 1.3312E-13 | 4.82943E-13 | Group07 | 3.410641193 | 50 |
| GO:0009653 | anatomical structure morphogenesis | 1.7424E-13 | 4.82943E-13 | Group07 | 2.48262167 | 75 |
| GO:0050793 | regulation of developmental process | 7.6418E-12 | 4.82943E-13 | Group07 | 2.409638643 | 68 |
| GO:0051239 | regulation of multicellular organismal process | 2.4253E-13 | 4.82943E-13 | Group07 | 2.421153307 | 76 |
| GO:0048731 | system development | 1.4322E-13 | 4.82943E-13 | Group07 | 2.065262318 | 100 |
| GO:2000026 | regulation of multicellular organismal development | 9.7336E-13 | 4.82943E-13 | Group07 | 3.126915932 | 51 |
| GO:0007166 | cell surface receptor signaling pathway | 5.9316E-07 | 4.82943E-13 | Group07 | 1.951703548 | 59 |
| GO:0072359 | circulatory system development | 4.5325E-16 | 4.82943E-13 | Group07 | 4.018912315 | 51 |
| GO:0006793 | phosphorus metabolic process | 3.1874E-10 | 3.59645E-12 | Group08 | 2.252559662 | 66 |
| GO:1901564 | organonitrogen compound metabolic process | 1.7291E-09 | 3.59645E-12 | Group08 | 1.662672281 | 111 |
| GO:0019538 | protein metabolic process | 1.9055E-06 | 3.59645E-12 | Group08 | 1.584228158 | 90 |
| GO:0006796 | phosphate-containing compound metabolic process | 1.7569E-09 | 3.59645E-12 | Group08 | 2.201582432 | 64 |
| GO:0035556 | intracellular signal transduction | 1.7861E-09 | 3.59645E-12 | Group08 | 2.262611389 | 61 |
| GO:0043412 | macromolecule modification | 0.00034824 | 3.59645E-12 | Group08 | 1.5625 | 62 |
| GO:0036211 | protein modification process | 0.00052832 | 3.59645E-12 | Group08 | 1.562086344 | 59 |
| GO:0051246 | regulation of protein metabolic process | 5.5454E-10 | 3.59645E-12 | Group08 | 2.265486717 | 64 |
| GO:0050789 | regulation of biological process | 2.2564E-06 | 4.37614E-06 | Group09 | 1.259675264 | 166 |
| GO:0048518 | positive regulation of biological process | 4.5662E-13 | 4.37614E-06 | Group09 | 1.780761957 | 122 |

| | | | | | | |
|---|---|---|---|---|---|---|
| GO:0051716 | cellular response to stimulus | 1.7638E-05 | 4.37614E-06 | Group09 | 1.370483637 | 121 |
| GO:0019222 | regulation of metabolic process | 3.183E-07 | 4.37614E-06 | Group09 | 1.530254126 | 109 |
| GO:0050794 | regulation of cellular process | 7.5248E-05 | 4.37614E-06 | Group09 | 1.238439918 | 154 |
| GO:0009893 | positive regulation of metabolic process | 4.1427E-07 | 4.37614E-06 | Group09 | 1.78012991 | 74 |
| GO:0048522 | positive regulation of cellular process | 4.7995E-13 | 4.37614E-06 | Group09 | 1.853055954 | 114 |
| GO:0031323 | regulation of cellular metabolic process | 1.4543E-05 | 4.37614E-06 | Group09 | 1.515151501 | 89 |
| GO:0060255 | regulation of macromolecule metabolic process | 1.2741E-06 | 4.37614E-06 | Group09 | 1.531463265 | 101 |
| GO:0080090 | regulation of primary metabolic process | 3.2942E-06 | 4.37614E-06 | Group09 | 1.535477638 | 95 |
| GO:0010604 | positive regulation of macromolecule metabolic process | 6.7513E-07 | 4.37614E-06 | Group09 | 1.809124231 | 69 |
| GO:0031325 | positive regulation of cellular metabolic process | 6.6741E-06 | 4.37614E-06 | Group09 | 1.78997612 | 60 |
| GO:0048519 | negative regulation of biological process | 3.7555E-13 | 4.95275E-05 | Group10 | 1.895168185 | 111 |
| GO:0050794 | regulation of cellular process | 7.5248E-05 | 4.95275E-05 | Group10 | 1.238439918 | 154 |
| GO:0007165 | signal transduction | 0.00073093 | 4.95275E-05 | Group10 | 1.345480084 | 96 |
| GO:0009892 | negative regulation of metabolic process | 1.3017E-07 | 4.95275E-05 | Group10 | 1.96374619 | 65 |
| GO:0048523 | negative regulation of cellular process | 1.9076E-12 | 4.95275E-05 | Group10 | 1.952822924 | 101 |
| GO:0031323 | regulation of cellular metabolic process | 1.4543E-05 | 4.95275E-05 | Group10 | 1.515151501 | 89 |
| GO:0051171 | regulation of nitrogen compound metabolic process | 2.8125E-06 | 4.95275E-05 | Group10 | 1.54922533 | 93 |
| GO:0060255 | regulation of macromolecule metabolic process | 1.2741E-06 | 4.95275E-05 | Group10 | 1.531463265 | 101 |
| GO:0080090 | regulation of primary metabolic process | 3.2942E-06 | 4.95275E-05 | Group10 | 1.535477638 | 95 |
| GO:0010605 | negative regulation of macromolecule metabolic process | 2.4074E-07 | 4.95275E-05 | Group10 | 1.989562988 | 61 |
| GO:0031324 | negative regulation of cellular metabolic process | 1.4402E-06 | 4.95275E-05 | Group10 | 2.043318272 | 50 |
| GO:0051172 | negative regulation of nitrogen compound metabolic process | 1.1699E-07 | 4.95275E-05 | Group10 | 2.113207579 | 56 |
| GO:0032501 | multicellular organismal process | 4.0236E-08 | 1.24474E-08 | Group11 | 1.461146832 | 132 |
| GO:0032502 | developmental process | 2.2457E-13 | 1.24474E-08 | Group11 | 1.785200119 | 124 |
| GO:0040011 | locomotion | 1.3312E-13 | 1.24474E-08 | Group11 | 3.410641193 | 50 |
| GO:0048856 | anatomical structure development | 1.9504E-14 | 1.24474E-08 | Group11 | 1.895066619 | 121 |
| GO:0048869 | cellular developmental process | 1.9361E-12 | 1.24474E-08 | Group11 | 2.000412464 | 97 |
| GO:0007275 | multicellular organism development | 6.7366E-14 | 1.24474E-08 | Group11 | 2.011656284 | 107 |
| GO:0009653 | anatomical structure morphogenesis | 1.7424E-13 | 1.24474E-08 | Group11 | 2.48262167 | 75 |
| GO:0050793 | regulation of developmental process | 7.6418E-12 | 1.24474E-08 | Group11 | 2.409638643 | 68 |

| | | | | | | |
|---|---|---|---|---|---|---|
| GO:0051239 | regulation of multicellular organismal process | 2.4253E-13 | 1.24474E-08 | Group11 | 2.421153307 | 76 |
| GO:0009888 | tissue development | 1.3806E-17 | 1.24474E-08 | Group11 | 3.209764957 | 71 |
| GO:0030154 | cell differentiation | 9.4839E-13 | 1.24474E-08 | Group11 | 2.017470837 | 97 |
| GO:0048513 | animal organ development | 2.2063E-13 | 1.24474E-08 | Group11 | 2.19644618 | 89 |
| GO:0048468 | cell development | 1.7538E-07 | 1.24474E-08 | Group11 | 2.10003829 | 55 |
| GO:0048731 | system development | 1.4322E-13 | 1.24474E-08 | Group11 | 2.065262318 | 100 |
| GO:2000026 | regulation of multicellular organismal development | 9.7336E-13 | 1.24474E-08 | Group11 | 3.126915932 | 51 |
| GO:0007399 | nervous system development | 1.4157E-07 | 1.24474E-08 | Group11 | 2.089552164 | 56 |
| GO:0072359 | circulatory system development | 4.5325E-16 | 1.24474E-08 | Group11 | 4.018912315 | 51 |
| GO:0002376 | immune system process | 0.00052624 | 0.000106114 | Group12 | 1.638291001 | 51 |
| GO:0023052 | signaling | 5.3299E-06 | 0.000106114 | Group12 | 1.443569541 | 110 |
| GO:0040011 | locomotion | 1.3312E-13 | 0.000106114 | Group12 | 3.410641193 | 50 |
| GO:0050896 | response to stimulus | 4.6119E-07 | 0.000106114 | Group12 | 1.367505074 | 143 |
| GO:0065007 | biological regulation | 2.701E-06 | 0.000106114 | Group12 | 1.236965179 | 172 |
| GO:0050789 | regulation of biological process | 2.2564E-06 | 0.000106114 | Group12 | 1.259675264 | 166 |
| GO:0007154 | cell communication | 1.473E-06 | 0.000106114 | Group12 | 1.468676925 | 113 |
| GO:0008219 | cell death | 1.0922E-10 | 0.000106114 | Group12 | 2.497883081 | 59 |
| GO:0009719 | response to endogenous stimulus | 6.8122E-14 | 0.000106114 | Group12 | 3.183627129 | 56 |
| GO:0042221 | response to chemical | 1.0697E-13 | 0.000106114 | Group12 | 1.930036187 | 112 |
| GO:0048518 | positive regulation of biological process | 4.5662E-13 | 0.000106114 | Group12 | 1.780761957 | 122 |
| GO:0048519 | negative regulation of biological process | 3.7555E-13 | 0.000106114 | Group12 | 1.895168185 | 111 |
| GO:0051716 | cellular response to stimulus | 1.7638E-05 | 0.000106114 | Group12 | 1.370483637 | 121 |
| GO:0023051 | regulation of signaling | 1.125E-10 | 0.000106114 | Group12 | 2.136031389 | 76 |
| GO:0048583 | regulation of response to stimulus | 1.3944E-07 | 0.000106114 | Group12 | 1.801169634 | 77 |
| GO:0050794 | regulation of cellular process | 7.5248E-05 | 0.000106114 | Group12 | 1.238439918 | 154 |
| GO:0007165 | signal transduction | 0.00073093 | 0.000106114 | Group12 | 1.345480084 | 96 |
| GO:0010033 | response to organic substance | 3.2473E-16 | 0.000106114 | Group12 | 2.477973461 | 90 |
| GO:0012501 | programmed cell death | 7.8421E-12 | 0.000106114 | Group12 | 2.680221796 | 58 |
| GO:0048522 | positive regulation of cellular process | 4.7995E-13 | 0.000106114 | Group12 | 1.853055954 | 114 |
| GO:0048523 | negative regulation of cellular process | 1.9076E-12 | 0.000106114 | Group12 | 1.952822924 | 101 |
| GO:0048584 | positive regulation of response to stimulus | 2.6757E-06 | 0.000106114 | Group12 | 2.004811525 | 50 |
| GO:0070887 | cellular response to chemical stimulus | 8.2297E-16 | 0.000106114 | Group12 | 2.523752928 | 85 |
| GO:1901700 | response to oxygen-containing compound | 1.2479E-13 | 0.000106114 | Group12 | 2.991886377 | 59 |
| GO:0010646 | regulation of cell communication | 6.7332E-10 | 0.000106114 | Group12 | 2.085094452 | 74 |
| GO:0010941 | regulation of cell death | 2.7226E-11 | 0.000106114 | Group12 | 2.76762414 | 53 |
| GO:0006915 | apoptotic process | 2.3613E-12 | 0.000106114 | Group12 | 2.763220549 | 58 |
| GO:0007166 | cell surface receptor signaling pathway | 5.9316E-07 | 0.000106114 | Group12 | 1.951703548 | 59 |

| | | | | | | |
|---|---|---|---|---|---|---|
| GO:0009966 | regulation of signal transduction | 2.9493E-07 | 0.000106114 | Group12 | 1.962677002 | 61 |
| GO:0035556 | intracellular signal transduction | 1.7861E-09 | 0.000106114 | Group12 | 2.262611389 | 61 |
| GO:0043412 | macromolecule modification | 0.00034824 | 0.000106114 | Group12 | 1.5625 | 62 |
| GO:0071310 | cellular response to organic substance | 2.218E-13 | 0.000106114 | Group12 | 2.600829363 | 69 |
| GO:0036211 | protein modification process | 0.00052832 | 0.000106114 | Group12 | 1.562086344 | 59 |
| GO:0008152 | metabolic process | 3.7563E-09 | 8.50935E-15 | Group13 | 1.376699328 | 160 |
| GO:0006807 | nitrogen compound metabolic process | 6.2275E-06 | 8.50935E-15 | Group13 | 1.345246434 | 134 |
| GO:0009058 | biosynthetic process | 1.5648E-08 | 8.50935E-15 | Group13 | 1.674958587 | 101 |
| GO:0044237 | cellular metabolic process | 5.0064E-08 | 8.50935E-15 | Group13 | 1.4224751 | 140 |
| GO:0044238 | primary metabolic process | 1.6619E-08 | 8.50935E-15 | Group13 | 1.404574394 | 148 |
| GO:0048518 | positive regulation of biological process | 4.5662E-13 | 8.50935E-15 | Group13 | 1.780761957 | 122 |
| GO:0048519 | negative regulation of biological process | 3.7555E-13 | 8.50935E-15 | Group13 | 1.895168185 | 111 |
| GO:0065009 | regulation of molecular function | 2.7266E-08 | 8.50935E-15 | Group13 | 2.037617445 | 65 |
| GO:0071704 | organic substance metabolic process | 7.803E-10 | 8.50935E-15 | Group13 | 1.407697797 | 158 |
| GO:0019222 | regulation of metabolic process | 3.183E-07 | 8.50935E-15 | Group13 | 1.530254126 | 109 |
| GO:0006793 | phosphorus metabolic process | 3.1874E-10 | 8.50935E-15 | Group13 | 2.252559662 | 66 |
| GO:0009892 | negative regulation of metabolic process | 1.3017E-07 | 8.50935E-15 | Group13 | 1.96374619 | 65 |
| GO:0009893 | positive regulation of metabolic process | 4.1427E-07 | 8.50935E-15 | Group13 | 1.78012991 | 74 |
| GO:0034641 | cellular nitrogen compound metabolic process | 0.01241812 | 8.50935E-15 | Group13 | 1.276207805 | 84 |
| GO:0043170 | macromolecule metabolic process | 0.00064861 | 8.50935E-15 | Group13 | 1.271230578 | 122 |
| GO:0044249 | cellular biosynthetic process | 1.0902E-06 | 8.50935E-15 | Group13 | 1.591375828 | 93 |
| GO:0046483 | heterocycle metabolic process | 0.02775608 | 8.50935E-15 | Group13 | 1.258931637 | 74 |
| GO:0048522 | positive regulation of cellular process | 4.7995E-13 | 8.50935E-15 | Group13 | 1.853055954 | 114 |
| GO:0048523 | negative regulation of cellular process | 1.9076E-12 | 8.50935E-15 | Group13 | 1.952822924 | 101 |
| GO:0050790 | regulation of catalytic activity | 6.5645E-08 | 8.50935E-15 | Group13 | 2.200083017 | 53 |
| GO:1901360 | organic cyclic compound metabolic process | 0.00131907 | 8.50935E-15 | Group13 | 1.369642258 | 85 |
| GO:1901564 | organonitrogen compound metabolic process | 1.7291E-09 | 8.50935E-15 | Group13 | 1.662672281 | 111 |
| GO:1901576 | organic substance biosynthetic process | 5.0345E-08 | 8.50935E-15 | Group13 | 1.65372932 | 98 |
| GO:0006139 | nucleobase-containing compound metabolic process | 0.03167148 | 8.50935E-15 | Group13 | 1.25698328 | 72 |
| GO:0009889 | regulation of biosynthetic process | 0.00139215 | 8.50935E-15 | Group13 | 1.464530945 | 64 |
| GO:0019538 | protein metabolic process | 1.9055E-06 | 8.50935E-15 | Group13 | 1.584228158 | 90 |
| GO:0031323 | regulation of cellular metabolic process | 1.4543E-05 | 8.50935E-15 | Group13 | 1.515151501 | 89 |

| | | | | | | |
|---|---|---|---|---|---|---|
| GO:0051171 | regulation of nitrogen compound metabolic process | 2.8125E-06 | 8.50935E-15 | Group13 | 1.54922533 | 93 |
| GO:0060255 | regulation of macromolecule metabolic process | 1.2741E-06 | 8.50935E-15 | Group13 | 1.531463265 | 101 |
| GO:0080090 | regulation of primary metabolic process | 3.2942E-06 | 8.50935E-15 | Group13 | 1.535477638 | 95 |
| GO:0006796 | phosphate-containing compound metabolic process | 1.7569E-09 | 8.50935E-15 | Group13 | 2.201582432 | 64 |
| GO:0009059 | macromolecule biosynthetic process | 0.01224705 | 8.50935E-15 | Group13 | 1.339101791 | 65 |
| GO:0010467 | gene expression | 1.9559E-05 | 8.50935E-15 | Group13 | 1.490145802 | 93 |
| GO:0010604 | positive regulation of macromolecule metabolic process | 6.7513E-07 | 8.50935E-15 | Group13 | 1.809124231 | 69 |
| GO:0010605 | negative regulation of macromolecule metabolic process | 2.4074E-07 | 8.50935E-15 | Group13 | 1.989562988 | 61 |
| GO:0018130 | heterocycle biosynthetic process | 0.04720541 | 8.50935E-15 | Group13 | 1.296207428 | 54 |
| GO:0019438 | aromatic compound biosynthetic process | 0.0470976 | 8.50935E-15 | Group13 | 1.293103456 | 54 |
| GO:0031324 | negative regulation of cellular metabolic process | 1.4402E-06 | 8.50935E-15 | Group13 | 2.043318272 | 50 |
| GO:0031325 | positive regulation of cellular metabolic process | 6.6741E-06 | 8.50935E-15 | Group13 | 1.78997612 | 60 |
| GO:0043412 | macromolecule modification | 0.00034824 | 8.50935E-15 | Group13 | 1.5625 | 62 |
| GO:0044271 | cellular nitrogen compound biosynthetic process | 0.01003173 | 8.50935E-15 | Group13 | 1.348314643 | 66 |
| GO:0051172 | negative regulation of nitrogen compound metabolic process | 1.1699E-07 | 8.50935E-15 | Group13 | 2.113207579 | 56 |
| GO:0051173 | positive regulation of nitrogen compound metabolic process | 4.3244E-06 | 8.50935E-15 | Group13 | 1.810088992 | 61 |
| GO:1901362 | organic cyclic compound biosynthetic process | 0.00073616 | 8.50935E-15 | Group13 | 1.500115395 | 65 |
| GO:0010468 | regulation of gene expression | 5.1807E-05 | 8.50935E-15 | Group13 | 1.528637767 | 79 |
| GO:0010556 | regulation of macromolecule biosynthetic process | 0.00999595 | 8.50935E-15 | Group13 | 1.393302321 | 57 |
| GO:0019219 | regulation of nucleobase-containing compound metabolic process | 0.0264968 | 8.50935E-15 | Group13 | 1.333650827 | 56 |
| GO:0031326 | regulation of cellular biosynthetic process | 0.00131841 | 8.50935E-15 | Group13 | 1.470588207 | 63 |
| GO:0034654 | nucleobase-containing compound biosynthetic process | 0.0465677 | 8.50935E-15 | Group13 | 1.295210123 | 53 |
| GO:0036211 | protein modification process | 0.00052832 | 8.50935E-15 | Group13 | 1.562086344 | 59 |
| GO:0051246 | regulation of protein metabolic process | 5.5454E-10 | 8.50935E-15 | Group13 | 2.265486717 | 64 |
| GO:0051252 | regulation of RNA metabolic process | 0.02753374 | 8.50935E-15 | Group13 | 1.341589212 | 52 |

### Key resources table

| *REAGENT or RESOURCE* | *SOURCE* | *IDENTIFIER* |
|---|---|---|
| *Antibodies* | | |
| anti-FAK (clone 77) | BD Biosciences | Cat no. 610087; RRID: AB_397494 |
| anti-p130Cas (clone 21) | BD Biosciences | Cat no. 610271; RRID: AB_397666 |
| anti-GAPDH | Santa Cruz Biotechnology | Cat no. sc-25778; RRID: AB_10167668 |
| Chemicals, peptides, and recombinant proteins | | |
| Trizol | Invitrogen | Cat no. 15596018 |
| RNeasy kit | Qiagen | Cat no. 74106 |
| Experimental models: Cell lines | | |
| Mouse: WT MEFs | Gift from Dr. Richard Assoian, University of Pennsylvania (Philadelphia, PA, USA) | N/A |
| Oligonucleotides | | |
| siRNA targeting sequence: FAK #1: CCUAGCAGACUUUAACCAAtt | Ambion | ID: 157448 |
| siRNA targeting sequence: FAK #2: GGCAUGGAGAUGCUACUGAtt | Ambion | ID: 61352 |
| siRNA targeting sequence: p130Cas #1: GCCAAUCGGCAUCUUCCUUtt | Ambion | ID: 161328 |
| siRNA targeting sequence: p130Cas #1: GCUGAAACAGUUUGAGCGAtt | Ambion | ID: 161329 |
| Silencer negative control siRNA | Ambion | AM4611 |
| Software and algorithms | | |
| FastQC 0.11.9 | Babraham Bioinformatics [54] | https://www.bioinformatics.babraham.ac.uk/projects/fastqc/ |
| HISAT2 2.2.1 | Kim et al., 2015 [57] | https://github.com/DaehwanKimLab/hisat2 |
| MultiQC 1.9 | Ewels et al., 2016 [56] | https://github.com/MultiQC/MultiQC |
| featureCounts 2.0.0 | Liao et al., 2014 [58] | https://subread.sourceforge.net/featureCounts.html |
| DESeq2 1.42.0 R package | Love et al., 2014 [61] | https://github.com/thelovelab/DESeq2 |
| ggplot2 3.4.4 R package | Wickman et al., 2016 [59] | https://github.com/tidyverse/ggplot2 |
| pheatmap 1.2 R package | Kolde et al., 2012 [60] | https://github.com/raivokolde/pheatmap |
| g:Profiler | Reimand et al., 2007 [63] | https://biit.cs.ut.ee/gprofiler/gost |
| QIAGEN Ingenuity Pathway Analysis | Krämer et al., 2014 [64] | https://www.qiagenbioinformatics.com/products/ingenuity-pathway-analysis |
| Cytoscape 3.10.1 | Shannon et al., 2003 [84] | https://cytoscape.org/ |
| Transite 1.3.0 | Krismer et al., 2020 [68] | https://transite.mit.edu/ |
